# Distinct genetic architectures of mRNA expression and splicing QTL in the Diversity Outbred mouse population

**DOI:** 10.64898/2026.03.06.710203

**Authors:** Charles I Opara, Kelly A Mitok, Christopher H Emfinger, Katheryn L Schueler, Donnie S Stapleton, Nancy A Benkusky, Udaya Gadiparthi, Kalynn H Willis, Victor Ruotti, Brian S Yandell, Mark P Keller, Alan D Attie

## Abstract

Genetic association studies of mRNA abundance phenotypes link regulatory gene loci to mRNA abundance (quantitative trait loci; QTL). Here, we used liver RNA-seq data from a large cohort of the Diversity Outbred (DO) mouse population to map QTL for gene abundance (eQTL), isoform abundance (isoQTL), isoform ratios (irQTL), and splicing (sQTL). QTL studies using this heterogeneous mouse population have been limited to gene-level abundance of mRNAs and have not focused on mRNA isoforms or splicing. We show for the first time that genetic effects on expression are distinct from splicing in the DO mouse population. Using allele-effect patterns from local QTL for protein-coding gene-isoform pairs, we show that genetic variation drives allele-specific isoform usage, generating isoforms whose genetic signals diverge from their aggregated gene-level effects. We conducted pathway enrichment on distal eQTL and isoQTL hotspots and uncovered pathways not detected with eQTL. We then applied a composite mediation approach at these distal hotspots that compares gene-gene, isoform-isoform, and isoform-gene mediator models. By contrasting these causal models of transcriptional regulation, we identified unique associations between mRNA isoforms. We identified candidate genes at irQTL and sQTL hotspots that regulate mRNAs primarily through isoform usage and splicing, including *Alkbh1* and *Dicer1*. For these driver genes, we nominated novel candidate targets through mediation analysis and pathway associations with known RNA-binding targets. We integrated our QTL data with human genetic data, prioritizing effector genes in loci associated with metabolically relevant traits. Our data also suggest that sex and diet influence eQTL and isoQTL distinctly and primarily through distal-acting gene loci. Overall, our findings highlight distinctive genetic effects on transcriptional and post-transcriptional mechanisms of gene regulation.

**Author summary:** Quantitative trait loci (QTL) analysis allows us to identify variable regions of the genome that are associated with phenotypes. Quantitative measures of gene expression constitute the most common form of mRNA abundance phenotype. QTL analysis of gene expression (eQTL) has been applied extensively to identify loci associated with gene regulation in humans and model systems. However, gene regulatory mechanisms can be altered transcriptionally or post-transcriptionally via alternative mRNA splicing. eQTL studies capture mostly the effects of genetic variants on transcription, much less on post-transcriptional modifications. In this study, we aimed to characterize the effects of genetic variation on post-transcriptional mechanisms in the Diversity Outbred mouse population. This mouse model system has substantial genetic diversity comparable to the human population, which allows for translatable studies. Studies of genetic effects on gene expression have been extensively reported in this mouse population, but not on post-transcriptional regulation. Our data highlight the substantial effects of genetic variation on post-transcriptional mechanisms, distinct from transcription, in this mouse population.

## Introduction

Genetic association studies link DNA variants to physiological and molecular trait outcomes (1). When we use mRNA expression as a quantitative trait, we can identify DNA variants that regulate mRNA abundance. These are termed expression quantitative trait loci (eQTL). eQTL studies conventionally prioritize gene-level estimates of mRNA expression. This approach overlooks a major source of transcriptomic complexity in eukaryotes: alternative splicing. Alternative pre-mRNA splicing (AS) mediates the production of multiple distinct mRNA isoforms from a single gene, which can differ in structure and function (2,3). AS has been established as a key mechanism through which genetic variation mediates complex trait outcomes (4–6). Variations in isoform transcript structure and function provide an opportunity to discover genetic variants controlling alternative splicing, thus distinguishing between the genetic regulation of transcriptional and post-transcriptional mechanisms.

Measurement of transcript abundance (7–10) and AS events (11–15) is commonly used to study the genetic control of mRNA splicing. While transcript-based methods focus on identifying genetic regulation of isoform abundance (isoQTL) or isoform ratios (irQTL), event-based methods prioritize variants that directly impact alternatively spliced junctions (sQTL). Each approach has distinct advantages. Isoform-level metrics provide a direct measure of transcript diversity, while event-based methods are more sensitive to post-transcriptional regulation. Consequently, it can be beneficial to combine quantification strategies to allow for the discovery of loci that may contribute to complex trait genes via different types of post-transcriptional regulation.

Advances in sequencing and analytical methods have enabled large-scale genome-wide association studies (GWAS) in both humans and model organisms. However, our ability to study genetic mechanisms underlying complex traits in humans remains limited by the practical challenges of establishing well-controlled, long-term, context-dependent studies. This necessitates the use of genomic approaches that prioritize model organisms and allow for translatable findings in humans.

We have adopted the Diversity Outbred (DO) mouse stock, originating from 8 inbred mouse strains (16), as a model system to study the genetic regulation of gene expression. Because of meiotic recombination during outbreeding, each DO mouse is genetically unique, and together, the DO mice offer the same level of genetic diversity as the entire human population (∼40 million SNPs). With a large sample size, we associate physiological and molecular phenotypes with variable regions of the genome (17,18). This approach is based on the premise that key metabolic pathways and gene regulatory mechanisms are highly conserved between mice and humans (19–21). As a result, the identification of these genetic mechanisms in a mouse population allows for the translation of physiological outcomes to humans.

Each DO mouse inherits a unique combination of the eight founder alleles in small haplotype blocks, and we can test whether differences in these haplotypes are associated with differences in the expression of a given mRNA across the population. For each significant locus, the effects of the eight founder alleles can be estimated to identify which founder haplotypes are associated with increased or decreased expression of the mRNA. These founder allele effect patterns capture the genetic architecture underlying mRNA expression and represent the basis of genetic mapping using this mouse population (18). Genetic effects on expression traits in this mouse population have been well established. Previous studies have investigated eQTL in islets (22), heart (23,24), brain (25), skeletal muscle (26), kidney (27), embryonic stem cells (28), and liver (29–33).

QTL studies on post-transcriptional regulatory mechanisms have been relatively unexplored in the DO mouse population. In this study, we utilized a large DO mouse cohort consisting of both male and female mice, fed one of two extreme diets: a low-fat/high-carbohydrate (HC) diet or a low-carbohydrate/high-fat (HF) diet. We performed QTL mapping to identify loci associated with RNA phenotypes at four different levels: eQTL, isoQTL, irQTL, and sQTL. We show that genetic effects on expression are distinct from splicing. Further, by comparing causal models of isoform abundance in addition to gene abundance, we identified unique associations between mRNAs not detected at the gene level. Notably, we identified candidate drivers that mediate complex trait mRNAs primarily through post-transcriptional regulatory mechanisms. By integrating human GWAS data, we show that candidate drivers of post-transcriptional regulation are associated with comparable traits in humans. Finally, we extend our analysis to show that sex and diet also influence the genetic regulation of mRNAs distinctly on eQTL and isoQTL.

## Results

### Genetic effects on transcriptional and post-transcriptional regulation in the DO mouse population

We performed RNA-seq on livers from 1,157 (male and female) DO mice maintained on one of two diets: a low-fat/high-carbohydrate (HC) diet or a low-carbohydrate/high-fat (HF) diet for 12 weeks (see Methods). This allowed us to quantify four levels of RNA molecular phenotypes: gene- and isoform-level mRNA expression, isoform ratios, and intron-excision ratios (**Fig 1A**). We identified ∼28,000 gene- and ∼85,000 isoform-level mRNAs and ∼160,000 intron-excision ratios (**S1A** and **S1B Fig**). We then performed whole-genome scans using diet and sex as additive covariates, initially negating their influence, to identify ∼26,000 eQTL, ∼62,000 isoQTL, ∼52,000 irQTL, and 96,000 sQTL (FDR< 0.1; LOD ≥ 7.5; **Fig 1B**, **S1C Fig** and **S1-S4 Tables**).

**Fig 1.**
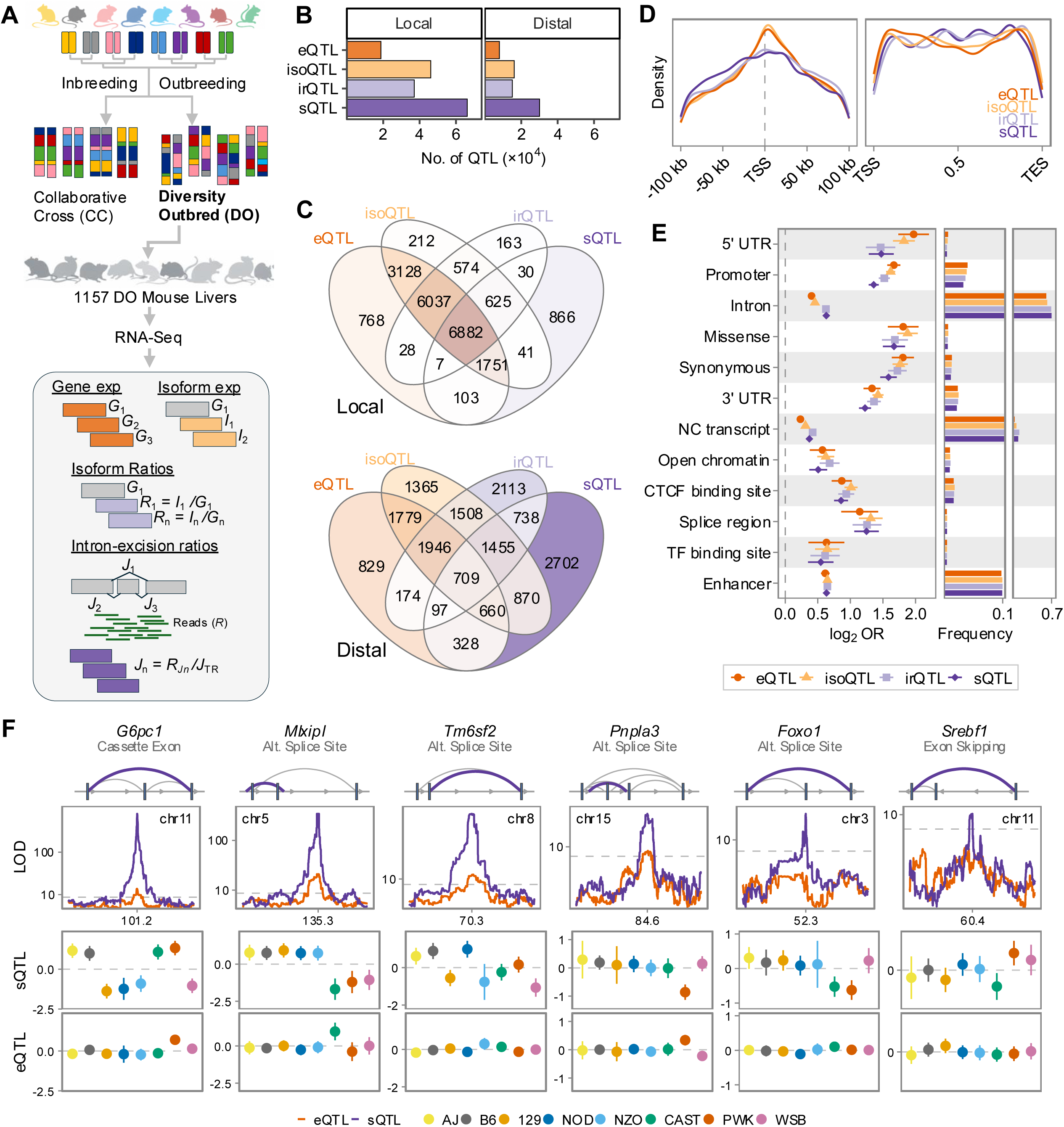
Genetic variation exerts distinct effects on post-transcriptional regulation in the DO mouse population. **(A)** Study Overview. Tissue samples from the livers of 1157 male and female mice of the Diversity Outbred (DO) mouse cohort were collected to quantify four molecular phenotypes: gene and isoform expression, isoform and intron-excision ratios, and subsequently perform QTL mapping. **(B)** Total number of local and distal QTL (FDR< 0.1; LOD ≥ 7.5) identified across molecular phenotypes. **(C)** Overlap of unique genes from all four molecular phenotypes for local and distal QTL, respectively. **(D)** Distributions of lead intragenic variants associated with local QTL around the transcription start site (TSS) (left) and relative to the gene structure (strand-adjusted and normalized to a median gene length of 48kb) (right). **(E)** Functional annotation of enrichment of lead intragenic variants associated with local QTL within a 3Mbp window (left) and frequency of the variants in each annotation class (right). For each consequence class, the enrichment of the lead variants was calculated relative to all tested variants in the same window (error bars indicate the 95% confidence interval). **(F)** Post-transcriptional regulation of key liver metabolic genes through genetic effects on mRNA splicing. For each gene, a schematic of the alternative splicing event (top row) associated with the splice junction (in purple), a comparison of the junction’s sQTL LOD profile and the gene’s eQTL (second row), and their corresponding allele effects (third and bottom row) is shown.

The majority (∼70%) of these QTL were characterized by local (within 4 Mb of the target gene TSS) regulatory elements. We found substantial overlap (∼32%) between the number of locally regulated unique genes identified from all four QTL analyses (**Fig 1C**). As expected, the majority (∼90%) of these were observed between eQTL and isoQTL. This observation, however, sharply contrasted with the marginal overlap (∼4%) between distally regulated unique mRNAs identified from our analyses. These results suggest that the majority of the local-acting genetic elements exert shared effects across the transcriptomic regulatory levels, while distal-acting elements exert distinct effects.

To further characterize these unique sets of mRNAs that correspond to the four QTL types, we examined the transcript annotations of unique genes and isoforms in our data (**S1D**-**S1F Fig**). Protein-coding mRNAs made up the majority of unique genes and isoforms in our data. Compared to distal, locally regulated isoforms were more frequently classified as alternate transcripts. We annotated splice junctions as constitutive if the start and end sites overlap an annotated intron boundary. We classified junctions overlapping one of two annotated intron boundaries as alternate. We classified junctions overlapping unannotated intron boundaries as novel/cryptic splice sites. Splice junctions overlapping annotated or one of the annotated splice sites constituted the majority of local sQTL **(S1H Fig)**. In many instances, multiple junctions overlapping annotated or unannotated splice sites belonged to the same gene, which is indicative of the effect of local genetic variants on alternative splicing mechanisms.

We next sought to distinguish between the effects of genetic variants on the RNA phenotypes in our study by examining single-nucleotide polymorphisms (SNPs) associated with the four types of QTL in our data. We evaluated the distribution of the lead intragenic variants (variants localized within the gene body) in each analysis relative to the transcription start and end sites (TSS/TES) and the gene body **(Fig 1D)**. Variants associated with the four types of QTL were differentially distributed around the TSS and relative to the gene body. As expected, lead variants associated with eQTL and isoQTL were enriched around the TSS, whereas lead variants associated with splice-related phenotypes (irQTL and sQTL) were concentrated along the gene body. These observations were reproduced when we repeated our analysis focusing on unique mRNAs (6,882) with a local QTL across all four layers **(S1G Fig)**. Further, functional annotation enrichment revealed that lead genetic variants located near upstream regulatory regions and promoter sites were most likely to affect expression traits, while variants in non-coding regions were strongly associated with splice-related phenotypes **(Fig 1E)**.

We evaluated cases where the regulation of an mRNA is predominantly explained by genetic effects on splicing, but not expression. We identified several core metabolic genes with well-established roles in liver metabolism, including *G6pc1*, *Mlxipl*, *Tm6sf2*, *Pnpla3*, *Foxo1*, *Srebf1*, *Hao1*, *Cyp7a1,* and *Apoe*. For these genes, we observed markedly significant QTL associated with alternatively spliced regions in the mRNA, but not for their expression **(Fig 1F** and **S1I Fig).** This suggests that for these genes and many others, their regulation is primarily mediated through mRNA splicing. Taken together, these results demonstrate the distinct contribution of sQTL to complex trait variation in the DO mouse population.

### The genetic architecture of QTL underlying transcript abundance and isoform ratio

We next sought to explore the possible divergence between local eQTL and isoQTL, focusing on protein-coding mRNAs. Conventionally, gene-level estimates of mRNA abundance are computed by aggregating transcript-level measurements in the form of transcripts per million (TPM) for all isoforms derived from the same gene. To identify which transcripts demonstrate similar or divergent genetic architecture to their gene-level QTL, we first computed pairwise correlations between gene and isoform expression QTL allele effects (**Fig 2A** and **S2A Fig**). Among 22,862 gene-isoform pairs, we observed a range of correlation values from −1 to +1, with ∼70% of isoforms showing substantial concordance with their gene-level QTL. We repeated our analysis correlating irQTL with eQTL, which resulted in only 25% of concordant isoforms, with 56% and 17% of isoforms showing substantial discordant or inverted signatures, respectively. When extended to all isoforms, we observed an even distribution of coding transcripts across correlation bins **(S2B Fig)**.

**Fig 2.**
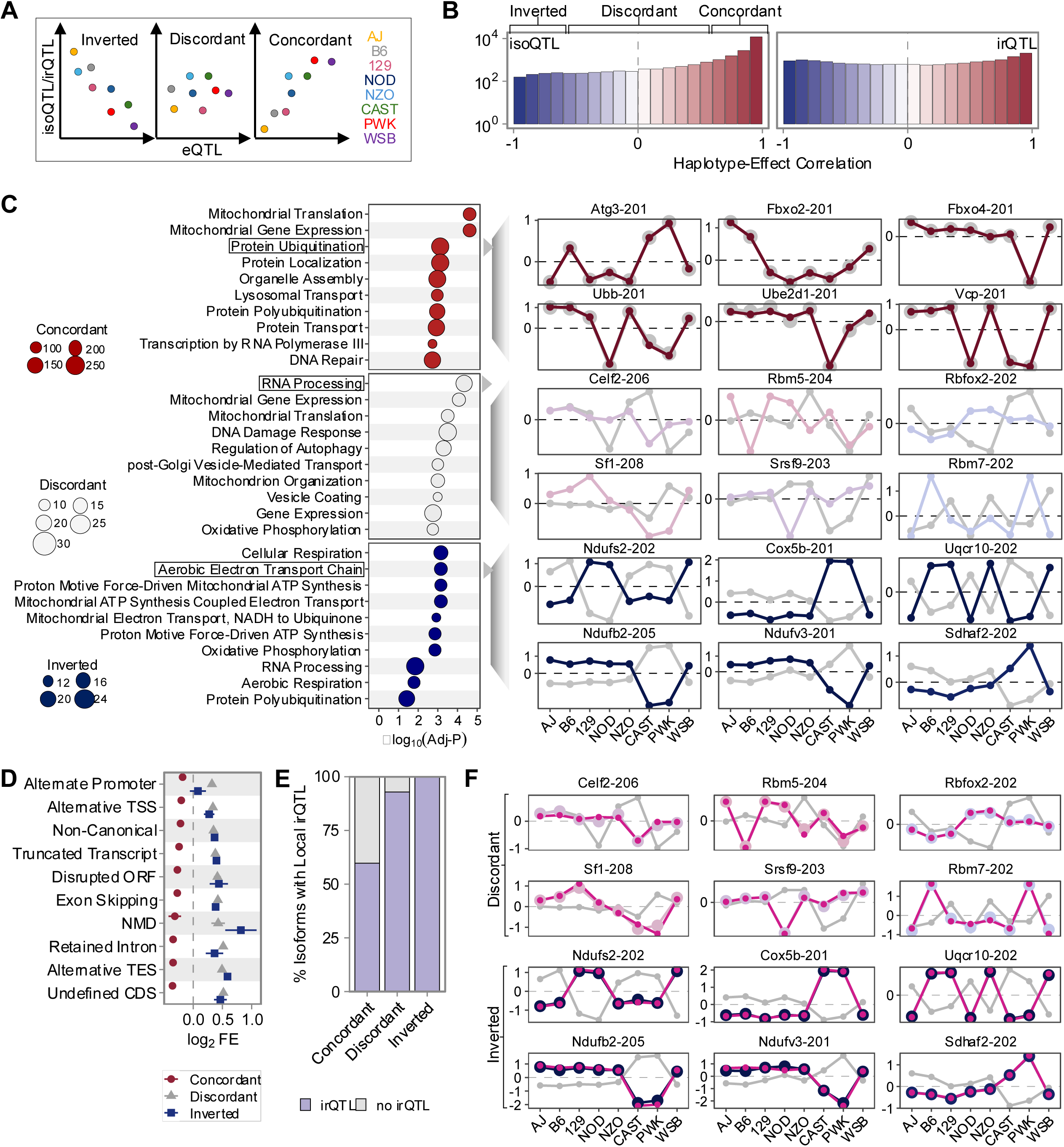
mRNA turnover underlies divergence of isoform QTL from their corresponding gene-level QTL. **(A)** Schematic of isoform classifications using allele effects from the eight founder strains, characterizing locally regulated isoQTL and irQTL to correlate with their corresponding local eQTL. Isoforms were classified into three categories: concordant (*r* ≥ 0.7), discordant (−0.7 < *r* < 0.7), or inverted (*r* ≤ −0.7) (See Methods). **(B)** The histogram shows the distribution of protein-coding isoforms captured in each bin based on the correlation coefficient with the local QTL allele effect and that of the corresponding gene QTL for isoQTL (n=22,862) (left) and irQTL (n=16,853) (right). **(C)** (Left) GO enrichment of isoQTL mRNAs across classes. Plots show the top ten enriched pathways (FDR < 0.05) across all classes (legends show the number of genes captured in the enriched pathways for each class). (Right) Representative examples of concordant, discordant, and inverted isoQTL relationships from genes captured in the highlighted pathways, showing allele effects when compared to their respective eQTL (eQTL allele effects are denoted by the grey dots/tracks). **(D)** Fold enrichment of genome-annotation features among isoforms from the three classes relative to all isoforms (*p* < 0.05). For each class and feature (GRCm39), significance was assessed by a one-sided Fisher’s exact test against all other matched isoforms. **(E)** Isoform usage characterizes discordant and inverted isoforms. Bar plots show the percentage of isoforms in each class that also had an irQTL. **(F)** Superimposed local irQTL allele effects (in pink) of the same discordant and inverted isoforms in **2C**.

The three correlation patterns (concordant, inverted, and discordant) between eQTL and isoQTL were enriched in distinct biological functions (**Fig 2C, S2C,** and **S5-S7 Tables**). The concordant mRNAs were especially enriched in those associated with translation and protein turnover. The discordant group contained mRNAs associated with translation and RNA splicing. The most homogeneous group was the one with the inverted pattern. It contained RNAs associated with mitochondrial function. In the case of transcript ratios, we observed no significant enrichments for biological functions, while the discordant and inverted transcript ratios showed no coherent enrichment structure **(S8-S9 Tables)**.

Notably, we observed that discordant and inverted isoQTL were far more likely to undergo post-transcriptional modifications compared to concordant isoforms **(Fig 2D)**, suggesting that these transcripts were more likely to be associated with genetic variants regulating post-transcriptional processes. To test this, we evaluated unique isoforms across these three classes, which are associated with both an isoQTL and an irQTL. We observed that discordant and inverted isoforms were more likely to have an irQTL compared to concordant isoforms **(Fig 2E)**. These discordant and inverted isoforms reproduced the same genetic architecture underlying their isoform ratios **(Fig 2F** and **S2D Fig)**. We observed less agreement with the sQTL signatures **(S2E Fig)**. These results distinguish genetic effects on transcript abundance from isoform usage. They also highlight genetic effects on isoform usage as a distinct regulatory mechanism that shapes overall transcript abundance.

### Causal modeling of gene and isoform-level associations with distal-mapping mRNAs

We have previously shown that many distally regulated QTL can co-map to the same locus (34,22). These “hotspots” often enrich for physiological function, suggesting a common regulator may mediate the expression of co-mapping mRNAs associated with a common pathway. We considered all distinct loci spanning an 8 Mbp window with ≥ 10 co-mapping distal QTL (**S3A Fig**). We further cataloged these into categories where eQTL co-map with or without the corresponding isoQTL. We identified hundreds of such loci, only a fraction of which were significantly enriched for biological function. In cases where eQTL and isoQTL co-map, these were relatively enriched for the same biological pathways (**Fig 3A**, **S10-S12 Tables**). Each of the hotspots was significantly enriched in one of several pathways (FDR<0.1), including translation, angiogenesis, mitochondrial respiration, immune response, transcriptional regulation, cell transport, lipid and carbohydrate metabolism, and protein metabolism (**S3B Fig**). We further observed that distal-mapping mRNAs uniquely detected for isoQTL were enriched for distinct biological pathways. These pathways were enriched in mRNAs associated with splicing, cellular transport, and protein metabolism.

**Fig 3.**
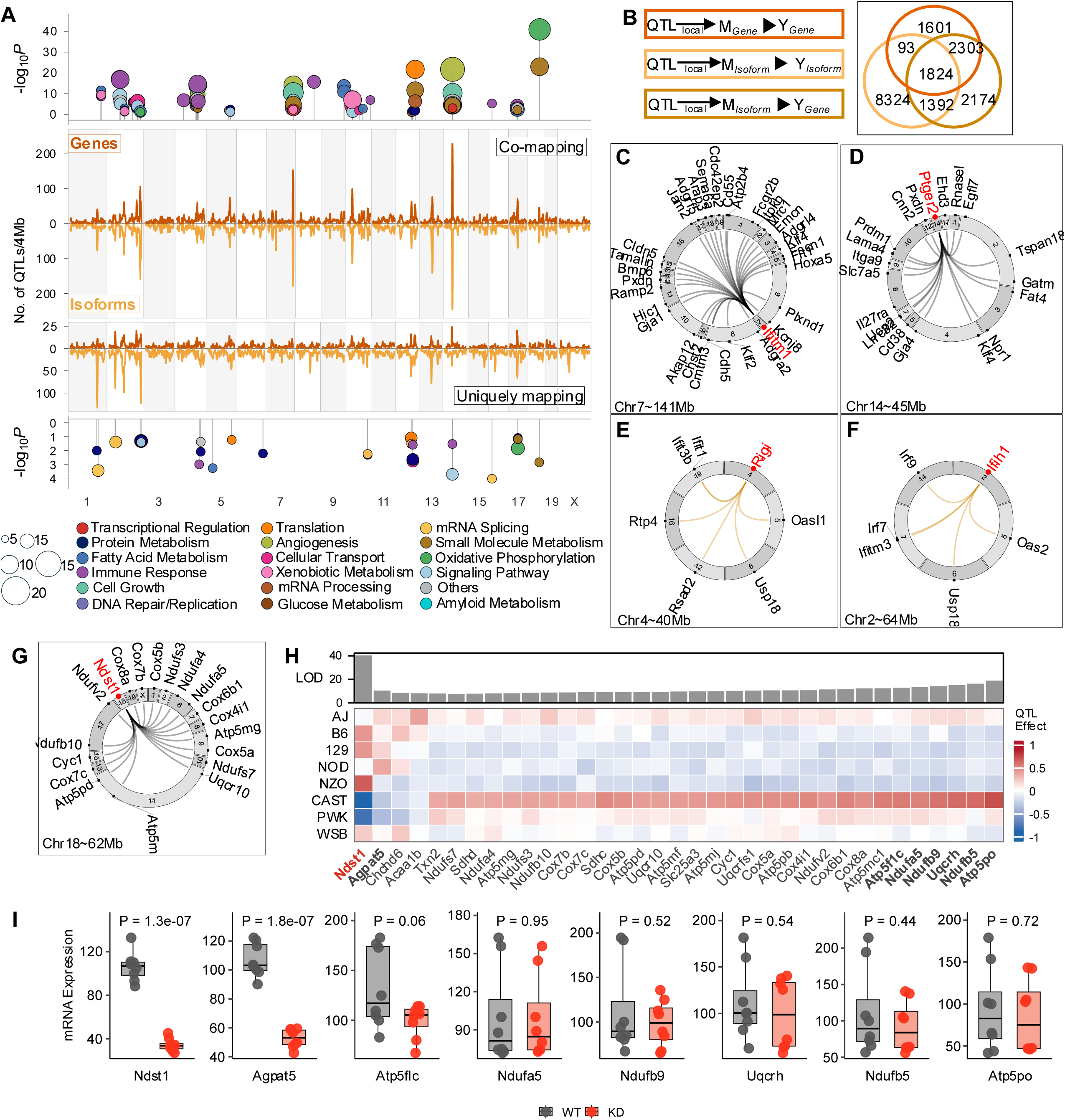
Distinct mediator-trait associations characterize gene- and isoform-level abundances. (**A**) Density plot showing the number of distal QTL (LOD ≥ 7.5; FDR < 0.1) occurring within a 4 Mbp window (middle panels) where eQTL and isoQTL co-map (top middle) or map distinctly (bottom middle). The top and bottom panels highlight pathway enrichment of the respective QTL loci. Each dot represents a unique set of genes (≥5) associated with the annotated pathways (FDR < 0.1). (**B**) (Left) Mediation analysis was performed using three scenarios for the identification of driver-trait relationships: (top) gene-level driver against gene-level mRNA abundance, (middle) isoform-level driver against isoform-level mRNA abundance, and (bottom) isoform-level driver against gene-level abundance. (Right) The Venn diagram shows overlaps in the number of unique gene-level collapsed mediator-trait pairs between models. (**C-F**) The circos plots show examples of mediator-trait relationships from all three mediation models at hotspot regions (inserted) that enrich for biological pathways in 3A (mediator genes (red), target genes captured in annotated pathway (black); connecting lines represent mapping of trait from genomic origin to distally located QTL; line colors represent mediator-trait relationships observed at both gene and isoform levels (black) or isoform-specific relationships (orange). (**G**) The circos plot shows *Ndst1* as a candidate driver of genes involved in oxidative phosphorylation at the hotspot on chromosome 18 (62Mb). (**H**) The heatmap shows QTL effects of *Ndst1* and the target genes (highlighted names represent genes whose expression was tested in *Ndst1* knockdown experiments). (**I**) The boxplots show mRNA expression of target genes (highlighted on the heatmap), comparing wild-type and *Ndst1* knockdown cells (*Ndst1^KD^*) (*P-values* inserted).

We next sought to identify candidate drivers at the distally regulated QTL hotspots that were enriched for physiological function. We performed mediation analyses to identify candidate drivers using bmediatR (35). Briefly, bmediatR utilizes a Bayesian model selection approach to infer a causal structure relating to the effect of the genotype at a QTL (X) on a target phenotype (Y) through a candidate mediator (M). It provides an advantage over other inference methods in that it is able to distinguish between complete mediation, partial mediation, and co-local effects, thereby limiting the detection of spurious mediators. Importantly, while complete mediators capture the full effect of the mediator on the trait, thereby making them potentially causal, partial mediators are unable to fully account for the effect of the mediator on the trait.

For our analyses, we considered only complete mediators under three models. Under the first two models, we used gene- or isoform-level expression as the mediator of either gene-level or isoform-level RNAs (**Fig 3B**). For the third model, we utilized isoform-level expression as the mediator of gene-level expression as the trait. This third model was largely driven by our discovery that isoforms from the same gene can demonstrate distinct genetic architecture, suggesting they will perform differently during mediation. Our results show an improved detection of driver-trait relationships when considering isoforms as mediators.

In several instances, we identified a mediator of mRNAs that co-maps to a hotspot enriched in a particular pathway (**Fig 3C** and **3D**, **S3C** and **S3D**). Interferon-induced transmembrane protein 1 (*Ifitm1*) and prostaglandin E receptor 2 (*Ptger2*) were identified as mediators of the angiogenesis-related hotspots on chromosomes 7 (∼141 Mb) and 14 (∼45 Mb), respectively. Both mRNAs have well-established roles in the regulation of angiogenesis (36,37). On chromosome 12 (∼100 Mb), we identified the gene encoding the deubiquitinating enzyme Ataxin-3 (*Atxn3*) as the candidate mediator of mRNAs involved in translation that co-map to the locus (**S3E** and **S3F**). These hotspots highlight examples where we detected the same mediators at the gene and isoform levels.

Notably, we discovered several instances where hotspots that enrich for the same pathways at the gene and isoform levels yielded a complete mediator only at the isoform level. For example, the hotspot on chromosome 4 (∼40 Mb), which was enriched for immune response pathways, was observed at both the gene and isoform levels. However, we identified the retinoic acid-inducible gene-I (RIG-I; *Rigi*) as the complete mediator at this locus only at the isoform level (**Fig 3E; S3G Fig**). The role of *Rigi* in viral RNA sensing and transcriptional induction of innate immune response genes is well established (38–40). When we mediated isoform-level abundances against gene-level abundance of the mRNAs that co-map to the locus, we were able to verify *Rigi* as the most likely mediator. This was also the case for another hotspot, on chromosome 2 (∼64 Mb), where we identified interferon-induced helicase C domain 1 (*Ifih1*) as the complete mediator of immune response mRNAs that co-map to the locus only at the isoform level (**Fig 3F; S3H Fig**). We identified *Ifih1* as the complete mediator at this locus only when we performed mediation using isoform-level abundance.

To further characterize mediator-target relationships identified in our analysis, we queried our data against the STRING protein-protein interaction database (v12.0, mouse) (41). We identified more protein-interaction pairs that were unique to the isoform-level mediator-trait pairs relative to the gene-level pairs (**S4A-S4D Fig**). This result provides evidence of association between these isoform-unique mediator-target relationships. It also highlights examples of likely missed signals that may result when isoform-level associations are neglected. Further, our analysis takes advantage of the distinct genetic architecture that can occur among isoforms for the same gene, which allows for the identification of novel associations.

We identified N-deacetylase-N-sulfotransferase 1 (*Ndst1*) as a candidate mediator of mRNAs that co-mapped to a hotspot on chromosome 18 (60 Mb) (**Fig 3G** and **3H, S4E** and **S4F Fig**). These mRNAs were mostly comprised of protein-encoding genes that make up the mitochondrial respiratory chain complexes. This locus showed the greatest pathway enrichment of all hotspots, showing a strong association with oxidative phosphorylation pathways (*P*-value ∼ 7.36E-41). QTL effect patterns at this locus were strongly driven by the CAST allele and highlighted an inverse relationship between *Ndst1* expression and the target genes that co-map to the locus. To validate *Ndst1* as a driver of these mitochondrial transcripts, we performed a knockdown of *Ndst1* in the AML-12 hepatocyte cell line and measured the expression of several mRNAs that have a distal-QTL to this locus (see Methods). We achieved a knockdown of ∼75% for *Ndst1* expression; however, we detected no significant change in the expression of mitochondrial complex proteins (*Atp51c, Ndufa5, Ndufb9, Uqcrh, Ndufb5,* and *Atp5po*) that mapped to the locus with the strongest LOD scores (**Fig 3I**). However, we observed that *Ndst1* depletion coincided with reduced *Agpat5* expression. *Agpat5* catalyzes the conversion of lysophosphatidic acid (LPA) to phosphatidic acid in glycerophospholipid biosynthesis and is known to localize to the mitochondria (42). While our results found no significant effects of *Ndst1* knockdown on the mitochondrial complex mRNAs in the liver, it suggests a possible link between heparan sulfate modification and glycerophospholipid metabolism in the liver.

A limitation of the reliance on mRNA expression QTL for mediation analysis is that it will miss instances where a missense mutation in a gene for which there is no expression QTL is, in fact, causal for a phenotype. Therefore, we examined coding SNPs within the region of the chromosome 18 hotspot and identified three missense variants (rs249835031, rs37683112, and rs237159338) that are unique to CAST and present in highly conserved regions of the peroxisome proliferator-activated receptor gamma coactivator 1-beta (*Ppargc1b*) gene (**S4G-S4J Fig)**. *Ppargc1b* is well established for its role as a master regulator of cellular energy metabolism and mitochondrial biogenesis (43). We detected no significant QTL for *Ppargc1b* in our analyses, and its mRNA expression showed no significant correlation with the mitochondrial genes that co-mapped at the locus. Our results suggest that missense variants within *Ppargc1b* likely mediate the expression of the mitochondrial mRNAs.

### Genetic drivers of post-transcriptional regulation

We next turned to the effects of distally acting genetic variants on post-transcriptional mechanisms. For this, we evaluated hotspots across the genome where multiple distal irQTL and sQTL co-map. We verified that sQTL junctions co-mapping to the same loci were mostly a single junction from a given mRNA that overlapped an annotated intron (**Fig A**). In many cases, distal irQTL converge at hotspots that are separate from sQTL, which further suggests distinct regulatory mechanisms underlying transcript ratio and intron-excision ratios (**Fig 4B**). In addition, both categories of distal QTL did not significantly overlap with those of the expression traits, further distinguishing between the two levels of transcriptional regulation. We identified 28 and 50 irQTL and sQTL hotspots, respectively, that were significantly enriched for function (**S5A** and **S5B Fig**, **S13** and **S14 Tables**). For example, the sQTL hotspots on chromosomes 1 (155Mb), 4 (10Mb), and 15 (86Mb) (**S5B Fig**) were not as strongly enriched for pathways with expression traits. These examples suggest that candidate drivers mediate the regulation of the associated mRNAs only through post-transcriptional mechanisms.

**Fig 4.**
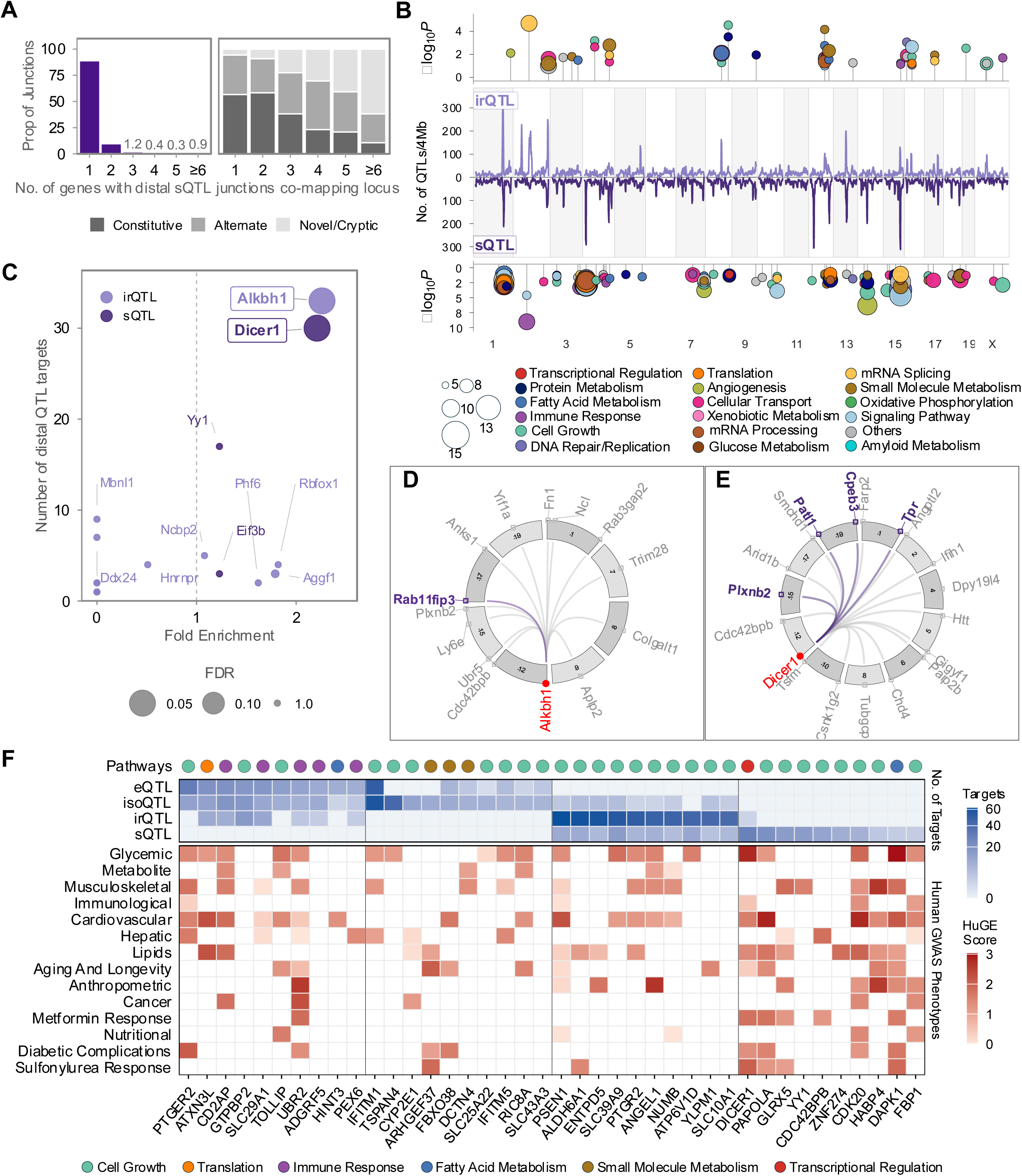
Candidate drivers of post-transcriptional regulatory mechanisms are associated with traits in human GWAS. **(A)** Bar plots show the proportion of splice junctions from a single gene that co-map to distinct distal QTL (left) and the junction annotation of junctions within each category (right). **(B)** Distal QTL mapping profiles of isoform- and intron-excision ratios per 4Mbp genomic window (LOD ≥ 7.5; FDR < 0.1) (middle panel). Pathway enrichments of the respective loci are indicated in the top and bottom panels. **(C)** Scatter plot shows the enrichment of RBPs relative to their number of distal sQTL targets. **(D-E)** The circos plots highlight known mRNAs with *Alkbh1* and *Dicer1* (in red) binding sites (in grey) and candidate targets captured in the same pathway with 2 or more of their known targets (in purple). **(F)** (Top) Circles indicate pathways associated with gene set targets of the drivers on the x-axis. (Middle) The heatmap shows the number of unique gene targets associated with the pathway categories above from all four QTL types. (Bottom) The heatmap shows the HuGE Score value for each driver gene associated with human GWAS categories.

A well-established mechanism by which genetic variants modulate post-transcriptional processing is through modifications of RNA-binding protein (RBP) sites. We performed mediation analysis by conditioning distal mapping irQTL and sQTL on isoform-level expression and prioritizing RBP isoforms. Our analysis identified 17 RBPs as mediators of irQTL and sQTL. To test these associations, we retrieved publicly available CLIP-seq datasets of RBPs from the ENCODE Project (44) and from the CLIPdb Database (45) (see Methods). We observed significant enrichment in binding sites associated with two RBP isoforms of the genes *Alkbh1* and *Dicer1*, for irQTL and sQTL targets, respectively (**Fig 4C**). 12 mRNAs out of the total 33 transcript targets contain binding sites associated with *Alkbh1*. The transcript isoform (*Alkbh1-201)* was the candidate mediator of transcript ratios from mRNAs including *Ntn1*, *Trim28*, *Fn1*, *Fgf1*, *Picalm*, *Plxnb2* and *Coro1b*, on chromosome 12 (∼87 Mb) (**S5C Fig**). These mRNAs were captured in overlapping pathways associated with the regulation of cell growth and projection. We identified *Dicer1-201* as the candidate mediator of sQTL from mRNAs on chromosome 12 (∼108 Mb) **(S5D Fig**). In the case of the *Dicer1* transcript, 14 out of 30 unique mRNAs associated with the corresponding distal sQTL are known binding targets of *Dicer1.* These were associated with multiple functions, including mRNA processing and stability, localization, and translation.

We next identified several additional candidate targets for these RBPs by incorporating mRNA targets that were captured in the same pathway with two or more of their known targets (**Fig 4D** and **Fig 4E**). For *Alkbh1*, two of its known targets (*Fn1* and *Plxnb2*) were captured in the same cell growth-related pathway as *Rab11fip3*. We also identified *Cpeb3*, *Patl1*, *Tpr,* and *Plxnb2* in various pathways with multiple known targets of *Dicer1*. In summary, our analysis not only identified *Alkbh1* and *Dicer1* as candidate drivers of distal irQTL and sQTL hotspots, but it also nominated additional target mRNAs that might be regulated by these RBPs.

We (22,46,47) and others (48–50) have shown that genetic associations identified in mice can be translated to humans. We next asked if the top candidate drivers across QTL types are associated with traits in humans. We identified human orthologs of the top candidate drivers across our QTL types and evaluated their loci for known associations with metabolic traits identified from human genome-wide association studies (GWAS) using the Human Genetic Evidence (HuGE) calculator (51). We observed substantial enrichments for candidate mediators at hotspots that regulate mRNA through splicing, compared to those that alter expression (**Fig 4F**). For example, the *Dicer1* locus in humans is strongly associated with glycemic traits and diabetes, which supports the roles of disease-associated genetic variants that modulate mRNA via alternative splicing. Further, it demonstrates our ability to recapitulate human-relevant metabolic signatures in our mouse population. Overall, our results demonstrate the significant role of post-transcriptional regulatory mechanisms underlying complex traits in the DO mouse population.

#### Sex and diet influence on gene and isoform expression QTL

We performed QTL analysis that allowed sex or diet to function as interacting variables (see Methods). This identified 7,869 sex-dependent and 4,424 diet-dependent eQTL and 18,218 sex-dependent and 11,478 diet-dependent isoQTL (FDR< 0.1; LOD ≥ 10.5; **S6A Fig, S15-S18 Tables**). However, in contrast to what we observed when sex and diet were additive covariates, ∼29% and ∼71% of both sex- and diet-dependent QTL were subject to local versus distal regulation, respectively. We observed more overlap between unique mRNAs that were influenced by sex and/or diet for local eQTL and isoQTL compared to distal (**S6B Fig**), suggesting that sex and diet exert isoform- specific effects not captured at the gene level. Pathway enrichment analysis identified a significant number of loci that enriched for biological functions at the isoform level, but not at the gene level (**S6C** and **S6D Figs; S19-S22 Tables**).

We observed a significant enrichment of mRNAs associated with cholesterol metabolism on chromosomes 7 (14 Mb) and 9 (121 MB) that are sex- and diet-dependent, respectively. A QTL that shows a strong sex or diet dependence suggests that the locus is affected by one sex or one diet. Our study consisted of 1,157 DO mice, half for each sex, and half for either the HC or HF diet (see Methods). When we stratified the mice by sex or diet, we detected distal eQTL and isoQTL present in one sex or diet but not the other (**S6E** and **S6F Figs**, **S23-S30 Tables**). This allowed us to ask which sex or diet was associated with the cholesterol biosynthesis pathways on chromosomes 7 and 9. We found that the sex-dependent hotspot on chromosome 7 (14 Mb) was specific to male mice (**S6G Fig**), while the diet-dependent hotspot on chromosome 9 (121 Mb) was specific to the HF diet (**S6H Fig**). The mRNAs that co-mapped to both loci encode enzymes that catalyze several core steps in the cholesterol biosynthesis pathway. QTL effect patterns at both sex- and diet-dependent loci were characterized mainly by the PWK strain.

We also observed substantial differences in the number of local mRNAs that were specific to either sex or diet (**S6I Fig**). For example, the stearoyl-CoA desaturase 1 (*Scd1*) and the patatin-like phospholipase domain-containing protein 3 (*Pnpla3*) local eQTL were only detected in one diet group (the HF diet group). These genes have been well studied in the context of diet-dependent regulation and liver disease development and progression (52–60), but not diet-dependent genetic association.

We evaluated human orthologs of our sex- and diet-specific mRNAs for known associations with metabolic traits in human GWAS. Our results highlight several mRNAs that were subject to sex- or diet-specific regulation in mice and have also been identified as effector genes within loci associated with the development or progression of metabolic phenotypes in humans (**S6J Fig**). For example, we identified several genes known to be influenced by oxidative, osmotic, and inflammatory response-induced changes in cellular environments. These changes are known to be sensitive to variations in nutrient availability within cells. We detected a significant HC-diet-specific signature associated with *Satb1* that suggests its role in regulating inflammatory response is also sensitive to nutrient composition. In humans, the *Satb1* gene locus is associated with several metabolic phenotypes, including nutritional and immune response-related traits. This result is relevant considering the reported association of the locus with inflammatory bowel disease in human colonoids (61). This finding suggests similar nutrient-sensitive roles for several other mRNAs, including the osmotic stress response genes serine protease 35 (*Prss35*) and the leucine-rich repeat-containing 8-E (*Lrrc8e*). Put together, our results show that context-specific genetic effects exert influence predominantly through distal-acting loci, and distinctly on gene and isoform mRNA abundance.

#### Interactive web resource for DO mouse QTL data

We provide an interactive QTL viewer (see Data) that enables users to easily query their gene of interest in our genetic data. This resource provides access to all genes and isoforms that exhibit genetic regulation in our DO dataset, allowing for a comparison of local and distal effects within a single, consistent workflow. To query an mRNA, users can enter a gene or transcript symbol into the search window, which returns a genome-wide LOD profile for the gene- or isoform-level trait, allowing users to immediately distinguish loci that regulate mRNA of interest. For each QTL peak exceeding genome-wide significance, the viewer provides allele-effect plots (founder haplotype effects), which provide the underlying genetic architecture at the locus. Queries can be performed for all QTL identified from our additive genome scans, as well as those that showed sex- and diet-dependence. For traits influenced by sex or diet, users can identify the specific sex or diet underlying the interactive signal. We also integrated the mediation analysis results for each mRNA. For a selected mRNA with a distal QTL peak, candidate mediators identified at the locus are shown with their probabilities of being a complete or partial driver of the trait. In sum, this resource allows users to extensively query the transcriptional regulatory landscape of mRNAs in the livers of our DO mouse cohort.

## Discussion

Genetic association studies have been applied to mRNA abundance to identify gene loci that control gene expression. We and others have carried out these studies in mouse intercrosses and, most recently, in the Diversity Outbred population. Studies on the transcriptomic regulatory landscape of the DO mouse population have generally not focused on genetic regulation of post-transcriptional mechanisms (62). Here, we present the first QTL mapping analyses distinguishing transcriptional and post-transcriptional regulation in the DO mouse liver. To the best of our knowledge, our study is the largest genetic screen of the hepatic transcriptome conducted in the DO mouse population.

Local genetic effects on mRNA expression and splicing traits were predominant in our study. In many cases, these QTL were shared across RNA phenotypes, indicating co-regulation of transcriptional and post-transcriptional mechanisms. This observation is consistent with previous reports of genetic effects on co-transcriptional regulation (63,15,64).

We, however, identified cases where genetic associations for gene expression were distinct from genetic regulation of RNA splicing. We found that variants associated with expression QTL were more clustered around the TSS, while variants associated with splicing phenotypes were more enriched within gene bodies. This finding is consistent with previous studies (4,8,65). Thus, genetic variation can affect gene regulation through mRNA splicing, contributing to complex traits (5).

QTL mapping in the DO population enables an estimation of the effect of each allele from each of the eight founder parent strains at each locus (18). These allele-effect patterns recapitulate the unit measures of the phenotype when centered at zero. Therefore, the estimated effect of each allele from the founder strains can be associated with an increase or a reduction in the expression of an mRNA at that locus. We identified differences in the local genetic architecture between genes and their transcript isoforms. These differences are attributable to diverging founder allele effects between genes and their transcript isoforms.

We identified three distinct patterns of local genetic variation, characterized by allele-effect relationships between genes and their transcript isoforms: concordant, inverted, or discordant. At the expression level, the majority of transcript isoforms showed agreement with their gene-level allele-effect patterns (concordant). This contrasted with the majority of transcript isoforms that were discordant for isoform usage. Isoform abundance QTL that were discordant or inverted were over-represented within transcription/splicing factors and mitochondrial complex proteins, respectively. These two classes of isoforms were mostly comprised of alternative or non-canonical isoforms. Importantly, we found that they were more likely to have shared QTL effects with their isoform usage QTL, suggesting that genetic effects on isoform usage shape overall transcript abundance. Our analysis here focused on protein-coding isoforms, which is consistent with previous studies that demonstrate irQTL mechanisms alter isoform usage and protein structure (8,66).

We identified several distal QTL hotspots where mRNAs detected at both the gene- and isoform-levels co-map to the same loci. These hotspots were enriched for the same biological pathways at both levels. However, when we performed mediation analysis at both levels, we identified several cases in which a causal mediator was identified only at the isoform level. This was the case for the hotspots on chromosomes 2 (64 Mb) and 4 (40 Mb), which were enriched in immune response mRNAs. Our approach to mediation prioritized the identification of potential causal mediators as complete if they capture the full effect of the mediator on the trait. From these examples, our results suggest that aggregating isoform-level abundance to gene-level estimates often masks the QTL effect of the causal transcript-isoform, resulting in weak or no associations.

Different isoforms produced from the same primary transcript often have different functions, including DNA and protein binding, transcriptional activation, and subcellular location. In one study (67), two-thirds of alternative TF isoforms exhibited significant differences in molecular interactions and functions, which classify them as negative regulators and rewirers. These variations in molecular interactions and functions of TF isoforms were attributed to differences in their sequences. We integrated our data with protein-protein interaction data to validate isoform-unique associations between mRNAs that were not detected at the gene level. We found several instances where the proteins of the mRNAs that were only associated at the isoform level were known to interact. Thus, inferring direct implications of our findings for the mouse hepatic interactome. Overall, these results show that prioritizing isoform-level quantification increases the discovery of trait associations (68).

We identified candidate mediators for all hotspots in our study, except the hotspot on chromosome 18 (60 Mb). Although our initial analysis flagged Ndst1 as the candidate mediator at the locus, experimental validation of the association proved otherwise. The potential pitfalls of genetic mediation analysis, as exemplified by this observation, have been addressed (69). The identification of Ndst1 as a driver at the locus resulted from the strong QTL signal and the matching allele effects between the mRNA’s local QTL and that of the co-mapping targets (the mitochondrial mRNAs). This highlights cases where a mediator is nominated at a QTL to be causal, when in fact the relationship is independent. In this case, we were able to rule out Ndst1 as the likely mediator based on our experimental data and the presence of another candidate mediator (Ppargc1b) within the QTL support interval, which had a greater biological plausibility.

We identified three coding variants in the Ppargc1b mRNA that were associated with the target mRNAs that co-mapped to the locus. Among the three coding variants that we identified in Ppargc1b, rs237159338 results in an amino acid change (Gly43Asp) that occurs in the N-terminal domain of the protein, which is critical for the transcriptional co-activator properties of this protein (70). This property of *Ppargc1b* is regulated via post-translational modifications (71). Under this mechanism, increased acetylation of amino acid residues in multiple domains of the protein represses its ability to co-activate transcription. Our findings therefore suggest that the variant alters the acetylation of amino acid residues in the N-terminal domain of the protein, thereby abolishing the repression of its activity and resulting in the increased expression of the mitochondrial complex proteins that co-map to the locus. This finding exemplified one of several instances where coding variants, associated with mRNAs that were undetected by mediation analysis, were the causal mediators of co-mapping distal traits. Together, our results highlight the likely limitations of the overreliance on mediation analysis alone when nominating mediators of traits in the DO mouse population.

We identified several candidate drivers of post-transcriptional mechanisms in our study. We found that *Alkbh1* mediated gene regulation through genetic effects on isoform usage, while *Dicer1* mediated gene regulation through regulation of splicing. These mRNAs are known to regulate RNA modifications through separate mechanisms.

The first mechanism involves RNA modifications via demethylation. *Alkbh1* belongs to the ALKBH class of proteins that encode enzymes that remove methyl groups occurring on N6 of adenosine (m^6^A) of mRNA molecules (72,73). These enzymes are known as demethylases and have been reported to regulate mRNA splicing, translation, and degradation (74–77). Under this mechanism, increased deposition of m^6^A on mRNAs results in reduced translation (78). Consequently, the activity of demethylases can be linked to post-transcriptional modifications and the subsequent utilization of their mRNA targets. Demethylases like *Alkbh1*, which target multiple RNA substrates, are often associated with diverse molecular functions. In our analysis, mRNA targets of *Alkbh1* included *Ntn1*, *Plxnb2*, *Fn1*, *Coro1b*, and *Picalm*. These are known binding targets of *Alkbh1* and were captured in shared pathways associated with the positive regulation of cell growth. This finding is consistent with the established role of *Alkbh1* in tumor and cancer progression (79–81). Our analysis also nominates *Rab11fip3* as a candidate target of *Alkbh1*. This mRNA was captured in the same cell growth pathways as the known targets of *Alkbh1* and has been implicated in cancer progression (82).

The second mechanism highlights post-transcriptional gene regulation mediated by microRNA (miRNA) biogenesis (83). *Dicer1* encodes an RNA endoribonuclease that cleaves pre-microRNAs (miRNAs) into mature miRNAs. These miRNAs are then loaded into the RNA-induced silencing complex (RISC) and control gene expression by hybridizing to target mRNAs and regulating their splicing, translation, or stability (84–88). This is in agreement with our finding of mRNA targets of *Dicer1* that were significantly enriched for mRNA processing and translation pathways in our analyses. We nominated several additional mRNAs associated with translation, including *Cpeb3*, *Patl1*, *Plxnb2*, and *Tpr* as candidate targets of *Dicer1*. Further, our results corroborate previous reports that found RBPs and miRNAs have stronger distal-acting effects on intron levels or exon-intron ratios (14,6,15).

Our analyses allowed us to nominate drivers of QTL in mice that were associated with related pathways in humans. Several mRNA targets of *Dicer1* were also captured in pathways significantly enriched in cell growth, signaling, and adhesion in our study. These pathways are known hallmarks of tumor growth. This result is also in line with known associations of *Dicer1* in cancer progression. Mutations of *Dicer1* have been shown to cause disruptions in post-transcriptional gene regulation and affect multiple cellular functions, particularly susceptibility to tumor growth (89–91). This characterizes a rare genetic condition in humans known as the DICER1 syndrome, in which cancer predisposition is caused by pathogenic variants in the gene (92–94). Further, our integration of human GWAS data highlighted associations of *Dicer1* with diabetes-related traits in humans. This observation is supported by findings that show beta-cell-specific deletion of *Dicer1* results in a disrupted miRNA network, impairment in insulin secretion, and glucose homeostasis, and diabetes development (95,96).

Sex and diet influenced the genetic regulation of mRNA expression primarily through distal-acting QTL in our study. We can attribute this to the effects of regulators such as transcription factors, RNA-binding proteins, and other signaling intermediates, whose downstream targets can be captured as distal-mapping mRNAs and further modulated by nutrient components or sex hormones (97,98). We identified several sexually dimorphic genes known to be influenced by hormonal signaling. For example, sex-specific expression of genes belonging to the sulfotransferase 2 (Sult2) family is well established (99–102). These enzymes are known to play crucial roles in drug metabolism and detoxification. We identified two members of this gene family (*Sult2a4* and *Sul2a6*) whose regulation was distinctly female-specific, consistent with previous findings. We also observed a markedly significant female-specific association of the prostaglandin E receptor 3 (*Ptger3*). In the liver, *Ptger3* is a well-established regulator of bile acid synthesis (103) and inflammation-driven tissue repair (104). We further identified metabolism-related genes, including the cholinergic receptor nicotinic alpha 2 subunit (*Chrna2*), the low-density lipoprotein (LDL) receptor-related protein 2 (*Lrp2*), and Caveolin 1 (*Cav1*), whose regulation was characterized by male-specific signatures. Further, our identification of a male- and an HF-diet-specific reduction in cholesterol biosynthesis mRNAs associated with the PWK allele is consistent with previous reports on the impact of sex and diet on liver transcriptional regulation (105).

Limited data on isoform-level transcriptome-wide association studies in the liver of humans necessitated that we focus our GWAS interrogation on locally regulated gene-level mRNAs. Nonetheless, we were able to identify sex- and diet-dependent mRNAs in our mouse population that were associated with metabolic traits in human GWAS. For example, mRNAs that were sex- or diet-dependent in the liver of our mice were strongly associated with glycemic traits in humans. This finding is in line with our previous report of concordance in diabetes-associated traits between mice and humans (22). We identified several mRNAs that exhibited sex- and diet-specific signatures and are situated in loci in humans with known metabolic phenotype associations. Our findings in mice suggest the sensitivity to sex bias and nutrient composition of these effector genes in genetically associated loci in humans. For example, inflammatory responses of the immune system can be elicited by the consumption of diets high in fats and sugars (106,107). These responses result in the activation of the Nuclear Factor kappa B (NF-kB) gene and subsequent production of pro-inflammatory cytokines that mediate a diverse set of targets (108). The regulation of the special AT-rich sequence binding protein 1 (*Satb1*) gene is mediated through a feedback mechanism involving NF-kB-cytokine (IL-4) signaling (109). *Satb1* encodes a transcription factor and is known to remodel chromatin and regulate T-cell development and function (110,111). Our data further nominate several candidate genes that are modulated in a sex- or diet-dependent manner in the development of metabolic phenotypes. Our analyses of sex and diet effects were restricted to expression traits only for practical reasons. Future work will be required to disentangle the effects of sex and diet on post-transcriptional mechanisms.

Isoform QTL mapping offers a more highly resolved view of transcript regulation compared to the standard gene-level estimation. However, several key issues limit our analyses. First, our estimation of isoform abundance and ratios in this study was derived from short-read RNASeq. Our analyses also relied on current transcript annotation quality, such that incomplete or inaccurate transcript annotations may obscure true isoform-specific effects or miss novel regulatory events. These limitations make full-length transcript reconstruction and accurate quantification difficult, particularly for complex genes and low-abundance isoforms. Secondly, we aligned gene and isoform RNASeq reads in this study to the reference mouse genome. Read alignment to individualized transcriptomes has been shown to improve gene and isoform abundance estimates (112). We mitigated this limitation by sequencing paired-end reads to a depth of 60 million, which resulted in a 98% read mapping rate. These limitations can be further resolved with complementary approaches, including improved annotation, long-read transcript profiling, functional prediction, and experimental validation of isoform-specific genetic effects.

In summary, our results demonstrate distinct genetic effects on transcriptomic regulatory mechanisms. We detail here for the first time the contribution of splice variants to complex trait genes in a DO mouse cohort. We report genetic associations between isoform-level RNAs that are undetected at the gene level. We identified candidate drivers of post-transcriptional mechanisms. We also demonstrate that our data allow for translation to humans by integrating human GWAS data. Our data also reveal that sex and diet influence the genetic regulation of mRNA expression distinctly at the gene and isoform levels, primarily through distal QTL. We provide access to these data through an interactive web resource.

## Methods

### Diversity Outbred mouse population and tissue collection

A total of 1177 male and female Diversity Outbred (DO) mice, across 7 generations (gens. 41-47), were obtained from The Jackson Laboratory at 4-5 weeks of age (J: DO, JAX stock #009376) and maintained in the AAALAC-accredited University of Wisconsin-Madison Department of Biochemistry animal vivarium. Over the course of approximately 2 years, DO mice were received in waves of 100 mice, 50 males and 50 females. 100 mice from litter 2 of the gen. 41, 100 mice from litter 1 of the gen. 42, and 100 mice from each of the litters 1 and 2 of generations 43-47 were obtained. Mice were maintained singly housed, on a 12-h light/12-h dark cycle (6 AM/6 PM), at a temperature range of 67-76 °C with a humidity range of 28-60% and had *ad libitum* access to food and water.

From arrival until 6 weeks of age, mice were fed a standard chow diet (Formulab Diet 5008, LabDiet). At 6 weeks of age, half of the mice (half of the females and half of the males) were switched to a custom-made low-fat, high-carbohydrate (HC) diet (10.4% kcal from fat, 75.5% kcal from carbohydrate, 14.2% kcal from protein, caloric density 3.6 kcal/g) (TD.200339, Envigo Teklad Diets) and the other half to a custom-made high-fat, low-carbohydrate (HF) diet (83.5% kcal from fat, 2.5% kcal from carbohydrate, 14.0% kcal from protein, caloric density 6.2 kcal/g) (TD.200625, Envigo Teklad Diets). Instead of cellulose, both diets contained a mix of 7 fiber sources: pectin, inulin, HI-MAIZE 260 (resistant starch), resistant wheat starch (Fibersym), fructooligosaccharide (FOS), beta-glucan, and glucomannan. The amount of fiber added to each diet was adjusted to be proportional to the diet’s caloric density, while the ratio of the different fibers relative to each other was kept consistent between the two diets. Body weight and food consumption were measured weekly (food consumption by weighing the amount of food placed in the hopper and that which remained each week).

Each wave of 100 mice was sacrificed within a 9-15-day span (median 10 days), at 18-19 weeks of age (12-13 weeks on diet). On the day of sacrifice, mice were fasted for 4 hours (5 AM-9 AM) and subjected to an intraperitoneal glucose tolerance test (ipGTT) (glucose dose 0.74g/kg lean body mass) with 6 blood samples taken over the course of 2 hours. After the last blood sample was taken, mice were sacrificed, and tissues were harvested and snap-frozen in liquid nitrogen before being stored at −80° °C. 23 out of the 1200 total mice required euthanasia before scheduled sacrifice for various unrelated health reasons, leaving 1177 mice for which tissues were collected. All mouse experiments were conducted in accordance with University of Wisconsin-Madison IACUC-approved protocols.

### Bulk RNA isolation and sequencing

Total RNA was isolated from a frozen piece of the left lateral liver lobe harvested from each DO mouse using the RNeasy Lipid Tissue Kit (QIAGEN) with on-column DNA digestion using the RNase-Free DNase Set (QIAGEN). RNA samples were then shipped to Novogene Corporation Inc. for sequencing.

### RNA sequence quantification and analysis

Messenger RNA was purified from total RNA using poly-T oligo-attached magnetic beads. After fragmentation, the first strand of cDNA was synthesized using random hexamer primers, followed by the second-strand cDNA synthesis using either dUTP for a directional library or dTTP for a non-directional library. For the non-directional library, it was ready after end repair, A-tailing, adapter ligation, size selection, amplification, and purification. The directional library was ready after end repair, A-tailing, adapter ligation, size selection, USER enzyme digestion, amplification, and purification. The library was assessed using Qubit and real-time PCR for quantification, and a Bioanalyzer for size distribution analysis. Quantified libraries were pooled and sequenced on Illumina platforms according to the effective library concentration and data amount to generate raw reads.

RNA-seq data in the form of FASTQ files were acquired from Novogene Inc. and were processed for all samples. We performed quality control checks on all FASTQ files using FastQC (113). We then aligned the preprocessed FASTQ files using the STAR alignment software (114) for all samples to the reference genome (GRCm39). Gene- and isoform-level quantifications (abundance and ratios) were performed using RSEM (115). Individual gene and isoform abundances were compiled using Tximport (116). Final raw counts were estimated as total counts per million (TPM) reads. We then retained only genes and isoforms with non-zero abundance values in ∼13% of all samples. TPM counts were then normalized by applying a variance-stabilizing transformation using the R DESeq2 package (117).

### Splice Junction Quantification

Intron-excision ratios were generated as percent spliced in (PSI) measures using

LeafCutter (118). We extracted splice junctions from RNA-seq preprocessed BAM files using Regtools (119). We then clustered the extracted splice junctions using the leafcutter_cluster_regtools.py script and default parameters. Intron clusters were then processed and normalized using the prepare_phenotype_table.py script to generate a splice junction phenotype matrix for sQTL mapping. We annotated junctions using the mouse Ensembl GRCm39 reference genome annotation.

### Mouse genotyping and genotype data generation

A tail clip was taken from each DO mouse upon arrival from The Jackson Laboratory. For the 1177 mice that completed the study, DNA was isolated from the tail clip and used for genotyping with the GigaMUGA (Neogen). Genotyping reports were organized and pre-processed for use with the R/qtl2 package (120) and the genome build GRCm39. Genotypes were encoded using directions and scripts from <u>Broman/qtl2</u>. Quality control was performed using directions and scripts from the genotype diagnostics for diversity outbred mice vignette at <u>Broman/qtl2</u>. The combined assessment of missing genotype calls, genotyping error rate, number of crossovers, and genotype frequencies for each sample was used to determine sample quality. Samples from 18 mice failed QC based on this assessment. Further, two samples were identified as duplicates (having matching genotype calls across the GigaMUGA). These 20 samples were excluded, leaving a total of 1157 samples (285 females on the HC diet, 286 females on the HF diet, 293 males on the HC diet, 293 males on the HF diet) for subsequent analyses. To validate the integrity of our remaining samples, we performed a concordance analysis between their genotype and expression data (121) (**S7A Fig**).

Genotyping markers were filtered to retain only those that were informative and of high quality. Those that did not uniquely map to the mouse genome or were monomorphic amongst the eight DO founder strains were removed. Furthermore, individuals with a large number of missing genotype calls, a high rate of genotyping errors, and/or unusual genotype frequencies were excluded. After this QC and filtering, 109,581 remained (109, 548 on chromosomes 1-19 and X) out of the 143,259 initial genotyping markers. Genotype probabilities, allele probabilities, and kinship matrices were calculated using the R/qtl2 package. Genotype probabilities were calculated using a hidden Markov model with an assumed genotyping error probability of 0.002, using the Haldane map function. Genotype probabilities were then reduced to allele probabilities, and allele probabilities were used to calculate kinship matrices, using the “leave one chromosome out” (LOCO) method.

### QTL Mapping

Normalized expression values for each expression trait were “RankZ”-transformed before QTL mapping. That is, expression values were converted to ranks, and then corresponding quantiles were found from the normal distribution. This is also known as “quantile-based inverse Normal transformation” or as transforming to “normal quantiles”. Genome scans were performed using a single-QTL linear mixed model with the scan1 function in the R/qtl2 package. Scans were carried out using data from the full set of 1157 mice, the allele probabilities, the kinship matrix calculated using the “leave one chromosome out” (LOCO) method, and with generation_litter (i.e., wave) and sex*diet as additive covariates. We included generation_litter as an additive covariate in our model as it accounted for substantial batch variation in our sample population (**S7B Fig**). Tissue harvesting and sequencing batches did not account for any variation in our samples.

We used permutation analysis to set the significance threshold for identifying significant QTL for our phenotypes following the steps described in (122). Briefly, RankZ-transformation of data gives a comparable null distribution per phenotype. This allows for the significance threshold that is representative of the full phenotype set to be obtained from a random subset of phenotypes. Calibrations of phenotype and permutation numbers found ∼100 phenotypes and ∼750 permutations optimal for determining the significance threshold for a standard QTL study. To determine the significance threshold for each data layer, we randomly sampled 100 gene-level phenotypes and 300 isoform and splice junction phenotypes. For each phenotype, we performed 1000 permutations using the scan1perm () function in the R/qtl2 package and calculated the significance threshold from each at a 10% significance level. At this significance threshold, we identified an LOD of 7.5 for all phenotype layers (**S1 Fig C**).

Peaks at or above the significance level were harvested, using the default setting of the find_peaks() function for allowing one peak per chromosome per trait. We then used the R/qvalue package (qvalue version 2.38.0) to first calculate the p-value for each peak LOD score from the gene (transcript) scans using the permutation LOD scores from the gene (transcript) scan, and subsequently to calculate the q-value for each peak. We controlled FDR at 10% across all phenotype peaks. We defined local-QTL as those located no more than 4 Mbp away from their respective gene’s start site, and all other QTL as distal-QTL. Estimated effect coefficients for the eight DO founder alleles were calculated at each peak using the fit1() function in the R/qtl2 package using the best linear unbiased prediction (BLUP) method. The genome scans described in this section are hereafter referred to as the “additive scans” in subsequent Methods sections.

### QTL x covariate interactions

We performed additional genome scans, permutations, and QTL peak identification on the expression traits exactly as described in the Methods section “QTL mapping”, except now with the addition of either sex or diet as an interactive covariate (“sex interactive” and “diet interactive” scans, respectively). As in the additive scans, we defined a significance threshold at a 10% significance level. This threshold was an LOD score of 10.5 for the permutation of either a sex interactive scan or a diet interactive scan, using either the expression data for a gene or for a transcript. We again controlled FDR at 10%. Local- and distal-QTL were defined as in the “QTL Mapping” Methods section. Estimated effect coefficients for the eight DO founder alleles at interactive scan peaks were calculated using the fit1() function but without using the BLUP method, as this method is not currently implemented to work with interactive covariates in R/qtl2.

To assess evidence for a QTL x sex or QTL x diet interaction, we considered significant QTL from the sex interactive and diet interactive scans, respectively, and calculated the difference between the LOD scores of these QTL and those at the same position from the additive scan for the same trait (regardless of whether or not the additive LOD score met the additive scan significance criteria). Importantly, before running each permutation analysis, the random number generator seed was set to the same predetermined value, ensuring that the 1000 permutation replicates were perfectly matched across all f permutations. This allowed us to calculate the difference between the LOD score of an interactive permutation and that of an additive permutation for each permutation replicate, and to use this set of permutation LOD score differences to establish a significance threshold for a significant QTL x covariate interaction. We defined a significance threshold at a 10% significance level, and this threshold was a LOD score difference of 4.1 for either a QTL x sex or QTL x diet interaction, using either expression data for a gene or isoform. In summary, we defined a QTL as one that met both our significance criteria for the relevant interactive scan and for the difference in interactive and additive LOD scores.

Genome scans were performed on subsets of the mouse cohort, stratified by sex or diet, to identify which sex or diet may be responsible for a significant QTL x sex or QTL x diet interaction identified in the genome scans using all mice, respectively. For these scans, the RankZ transformation was applied to the expression data after subsetting. Additive covariates used for scans of mice of one sex (one diet) were generation_litter and diet (generation_litter and sex). For each mouse set/covariate combination, we permuted one gene and one transcript scan 1000 times, computed the significance threshold at the 10% significance level for identifying significant peaks, and controlled the FDR at 10% as before.

### SNP association

We performed SNP association using the scan1snps function in the R/qtl2 package and following the workflow in the <u>R/qtl2 User Guide</u> for genome build GRCm39. A SNP or indel was considered for association analysis if it 1) was polymorphic amongst the DO founder strains, 2) had a genotype call in all DO founder strains and the genotype call was homozygous, and 3) had a high-confidence genotype call in at least 6 of the 8 DO founder strains. In brief, for each significant QTL peak, we imputed all SNPs and indels that met our criteria within a 3 Mbp window centered around each peak and performed a scan to test for association between each variant and the relevant expression trait, using the same covariates that produced the relevant QTL. Variants with the strongest associations were then identified by harvesting those variants with a LOD score within 1.5 LOD of that of the variant with the highest LOD score. Consequences for variants with the strongest associations were predicted using Ensembl VEP (version 114).

#### Variant consequence enrichment

For each local QTL associated with a phenotype, we considered the lead variant with the maximum LOD score, keeping the variant with the most severe function per QTL. For each consequence class, we computed enrichment among the lead SNPs relative to all variants tested in the same local windows (background: every high-confidence SNP, reduced to one variant per genotypically equivalent class and subsampled per window). For variants with multiple annotated functions, we collapsed to one class, prioritizing by Ensembl impact severity (donor/acceptor splice sites > coding > splice region > UTR > intron/non-coding > promoter). Enrichment of relevant genomic features, including splice donor/acceptor sites, intron/exon boundaries, promoter, enhancer, TF binding sites, CTCF binding sites, and open chromatin, was computed against the genome region annotation (GRCm39).

### Mediation analysis

For each significant distal-QTL from the additive scans, we identified those genes and transcripts with a significant additive local-QTL located within 4 Mbp of the distal-QTL. For each trait producing a local-QTL, we tested for mediation of the trait producing the distal-QTL using the R/bmediatR package (35). All expression data were RankZ-transformed before mediation analysis (see “QTL mapping” Methods section for an explanation of this transformation). Allele probabilities and the same covariates that produced the relevant QTL were used. All other arguments to the bmediatR () function were left at their defaults. We defined a mediator as having a posterior probability of mediation > 0.95 and a posterior probability of a co-local effect < 0.001. From there, each was labeled a “complete” mediator or “partial” mediator, depending on the mediation model that had the higher posterior probability (complete or partial).

### Correlation of gene- and isoform-level QTL allele effects

For each gene, isoforms with significant local QTL were paired to the corresponding gene-level QTL from the same gene. Gene-isoform pairs were retained only when the gene and isoform QTL peaks mapped to the same chromosome and were within ±4 Mb of one another; when multiple gene-level peaks satisfied this criterion for a given isoform, the nearest gene peak was selected (ties resolved by higher gene LOD). Founder allele effects were represented as vectors across the eight DO founder haplotypes (A-H). We then computed Pearson correlation coefficients between the isoform allele-effects and the gene allele-effects. Founders with missing allele-effect estimates in either vector were excluded from the calculation. *P*-values were calculated and adjusted for multiple testing using the Benjamini-Hochberg procedure. Correlation calculations were performed in R.

### GO analysis

Gene Ontology (GO) enrichment analysis of genes was performed using the R/clusterProfiler (123) package and R/Enrichr (124). GO terms with adjusted p-value less than 0.05 and gene sets of 5 or more mRNAs were considered significantly enriched.

### Enrichment of RBP Sites

We pooled CLIP datasets from mouse and human in CLIPdb (POSTAR3) and ENCODE eCLIP (HepG2 and K562 cells), mapping human genes to their respective mouse orthologs. Fold enrichment of RBP-bound targets among a candidate’s distal-mapping mRNAs was computed using the candidate’s full set of target genes as background. We determined a significance threshold by permuting 20,000 random target sets of equal size drawn from the background. Benjamini-Hochberg FDR correction was applied across the tested RBPs.

### Human GWAS integration

We obtained data on genes in loci associated with human GWAS traits from the Human Genetic Evidence portal available at <u>HuGE</u>. This resource integrates human genetic data from the Type 2 Diabetes Knowledge Portal (T2DKP) (125) to quantify the involvement of a gene in the development and progression of disease or phenotypes (51). We cross-referenced gene sets to identify their human orthologs in loci associated with one or more traits belonging to all phenotype categories in humans.

### Software and data availability

Raw sequence data are available at NCBI SRA under the project accession PRJNA1527584. Genotype data were deposited and are accessible at DOI: <u>Dryad Data</u>. An interactive web application accompanying this work, the AttieLab QTL Viewer, is freely available here: <u>Viewer</u>. Source code is available at <u>GitHub</u>. Additional information and archived versions are available here: https://doi.org/10.5281/zenodo.19961001.

### siRNA-mediated knockdown of Ndst1 in AML-12 hepatocytes

AML-12 mouse hepatocyte cells (ATCC CRL-2254) were maintained at 37°C in a humidified incubator with 5% CO_2_. Cells were cultured in DMEM/F-12 supplemented with 10% fetal bovine serum (FBS), 1× insulin-transferrin-selenium (ITS), and 40 ng/mL dexamethasone. Cells were passaged using 0.05% trypsin-EDTA and routinely monitored for contamination.

For gene knockdown experiments, AML-12 cells were plated in 12-well plates at 1.0 × 10D cells per well and allowed to adhere overnight to reach ∼40-50% confluency at the time of transfection. Cells were transfected with siRNA targeting Ndst1 (Ndst1*^KD^*) or a non-targeting scrambled siRNA control using Lipofectamine 3000 (Thermo Fisher Scientific), according to the manufacturer’s protocol, and harvested 24 h post-transfection for RNA extraction.

Total RNA was isolated using the RNeasy Plus Mini Kit (250) extraction kit from Qiagen (Hilden, Germany), following the manufacturer’s instructions. cDNA was synthesized from total RNA using the High-Capacity cDNA Reverse Transcription Kit from Applied Biosystems (Thermo Fisher Scientific), according to the manufacturer’s instructions.

Quantitative real-time PCR was performed on each sample and analyzed in technical replicates with primers from Integrated DNA Technologies (IDT, Coralville, IA, USA). Gene-specific reactions were normalized to B-actin. Experiments were performed with n=4 biological replicates per condition. Statistical analyses were conducted in GraphPad Prism. For comparisons between scram control and Ndst1*^KD^* groups, a two-tailed unpaired Student’s t-test was used to test for significance.

Primers:

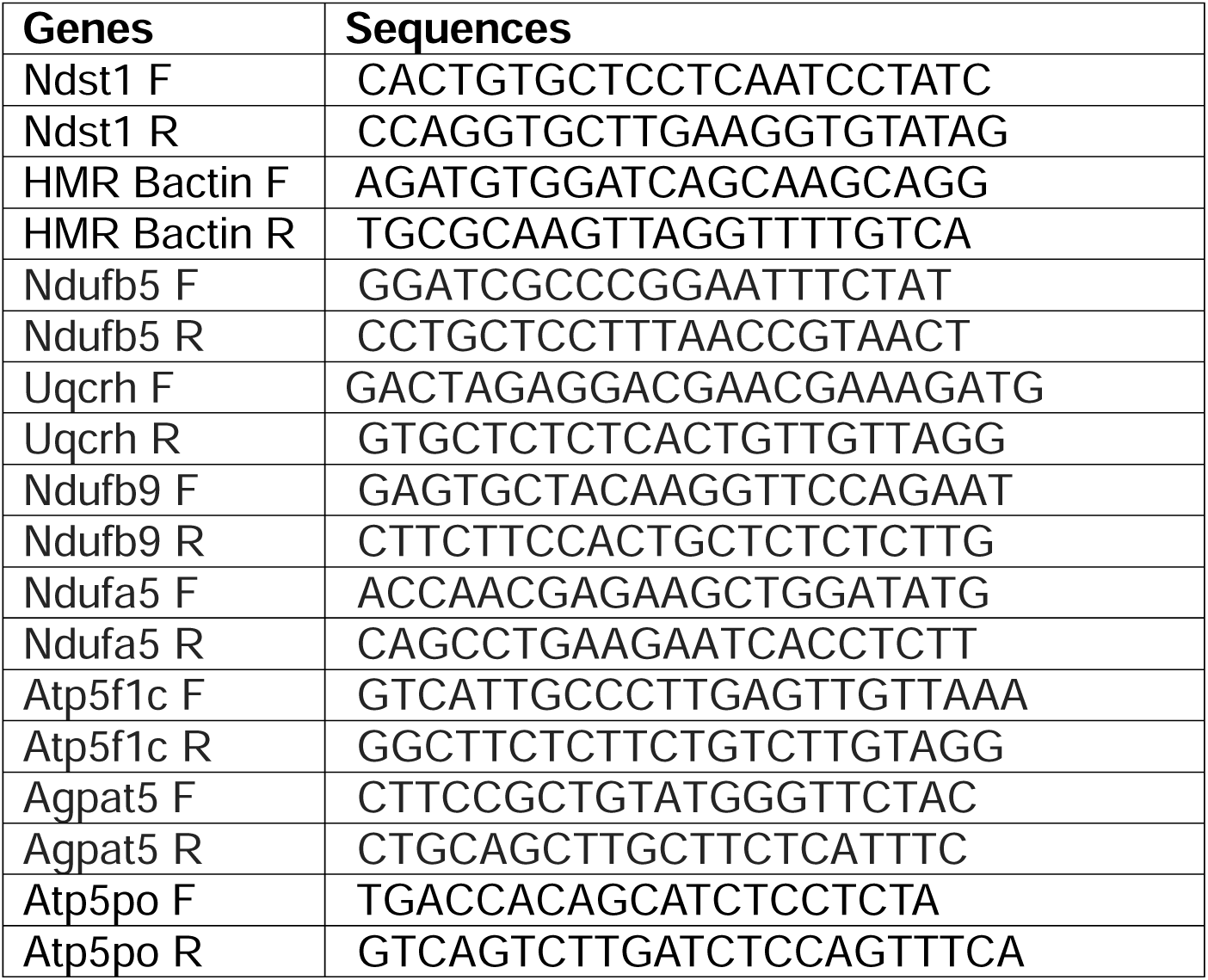

## Acknowledgments

We would like to thank Dr. Jason Flannick at the Broad Institute for providing access to the Human Genetic Evidence Data. This work was also made possible by computing resources at the Centre for High Throughput Computing and the Department of Biochemistry at the University of Wisconsin-Madison. This work was supported by funding from the National Institutes of Health grant (RC2DK125961).

## Supporting Information

### Supplementary Figures

**S1 Fig. (A)** Bar plot shows the number of quantified phenotypes per molecular feature layer. **(B)** Principal component analysis (PCA) of phenotype data. Plots show data for each molecular layer, before (top) and after covariate regression (bottom) (point shapes and colors represent sex and diet of each mouse sample). **(C)** Histograms show LOD distributions from permutations performed on a random set of phenotypes for each molecular trait layer. Permutations were performed using 100 gene-level expression traits, 300 isoform expression and ratio traits, and intron-excision traits. The dashed line indicates a ∼7.5 LOD score threshold that corresponds to an FDR < 0.1. **(D, E)** Transcript biotype composition of unique genes **(D)** and isoforms **(E)** associated with the various molecular QTL. **(F)** Transcript classification of unique isoforms corresponding to isoQTL and irQTL data. **(G)** Distributions of lead intragenic variants associated with local QTL, around the transcription start site (TSS) (left) and relative to the gene structure in mRNAs with a reported local eQTL, isoQTL, irQTL, and sQTL (normalized to a median gene length of 48kb) (right). Top panels display variants count and bottom panel show density normalized frequencies. **(H)** Splice site annotations of local sQTL junctions. For each annotation class (constitutive, alternate, novel/cryptic splice sites), the number of junctions associated with a unique gene is represented by the color shades. **(I)** Additional examples of key liver metabolic genes regulated primarily through splicing. Schematic of the alternative splicing event (top row) associated with the splice junction (in purple), a comparison of the junction’s sQTL LOD profile and the gene’s eQTL (second row), and their corresponding allele effects (third and bottom row) is shown.

**S2 Fig. (A)** Histogram distributions of isoform allele-effect correlation coefficients corresponding to isoform classes in **Fig 2B** (top) and threshold significance boundaries (bottom) indicate thresholds at |*r*| = 0.7. Because each correlation is computed across the same eight founders (*df* = 6), the significance cutoff follows the standard *t*-distribution mapping of r to its *p*-value from a two-sided *t*-test. At *p* < 0.05, this corresponds to 0.71. **(B)** Bar plots show transcript biotype compositions across the correlation coefficient bins corresponding to **Fig 2B**. **(C)** Scatter plots show the allele effect relationships between genes and isoforms in **Fig 2C**. **(D)** Allele effect correlations between overlapping isoQTL and irQTL isoforms. **(E)** Proportion of isoforms in each class with a corresponding sQTL (left) and allele effect correlation against their corresponding sQTL (right).

**S3 Fig. (A)** (Left) The number of hotspots (at a threshold of 10 co-mapping distal QTL within a 4Mbp window), as a function of co-mapping or distinct gene and isoform sets. (Right) The number of candidate hotspots compared to enriched hotspots across sets. **(B)** Gene Ontology Biological Process (GOBP) enrichment of selected hotspots. Pathways are shown as term-similarity networks, with nodes representing enriched terms. Encircled clusters represent enriched hotspots detected only uniquely mapping to distal isoform-level hotspots. Colors correspond to pathway categories in **Fig 3A**. **(C, D)** Heatmaps show allele effects of complete mediator-target relationships recovered at both gene and isoform levels: **(C)** *Ifitm1*; **(D)** *Pterg2*. (Mediator names are emboldened). **(E)** The circos plot shows *Atxn3* as a driver of the mRNAs associated with translation that co-map to chromosome 12 (100Mb), and **(F)** shows heatmaps of the allele effects of these co-mapping traits at both gene and isoform levels. **(G, H)** Heatmaps show allele effects of complete mediator-target relationships observed only at the isoform level: **(G)** *Rigi*; **(H)** *Ifih1*.

**S4 Fig. (A)** The bar plot shows the number of mediator-target interactions identified in the STRING (v12.0, mouse) protein-protein interaction network. For each mediation model, the number of pairs is shown as a function of interaction strength. (**B, C, D)** Interaction subnetworks show mediator-target relationships between protein-protein pairs from the isoform-isoform (mediator-target) unique pairs. **(E)** Stacked LOD profiles of the distal-mapping mitochondrial genes that co-map to the chromosome 18 (∼60 Mb) locus. **(F)** Mediation profiles of the top ten mitochondrial genes in **E**. For each target, the log posterior odds of mediation (top x-axis), posterior probability (top y-axis), and LOD drop fraction (bottom y-axis) are shown upon conditioning on *Ndst1* expression. **(G)** SNP profile shows CAST-private missense variants in the *Ppargc1b* gene locus associated with distal-mapping QTL of the mitochondrial genes. The highlighted red dots resolve to three unique lead variants in the gene’s locus: rs237159338, rs249835031, and rs37683112. **(H-J)** The plots show sequence alignment and conservation of all three SNPs: **(H)** rs237159338, **(I)** rs249835031, and **(J)** rs37683112 across model organisms.

**S5 Fig. (A)** (Left) The number of distal irQTL and sQTL hotspots (at a threshold of 10 co-mapping QTL within a 4Mbp window) and the number of candidate hotspots compared to enriched hotspots across sets (right). **(B)** GO enrichment of selected irQTL (top) and sQTL (bottom) hotspots. **(C)** Mediator-target allele heatmaps for *Alkbh1* (irQTL) and *Dicer1* (sQTL) primary transcripts. **(D)** Mediation profiles of select targets in **C**. For each target, the log posterior odds of mediation (top x-axis), posterior probability (top y-axis), and LOD drop fraction (bottom y-axis) are shown upon conditioning on transcript expression.

**S6 Fig. (A)** The bar plots show the total number of local and distal QTL significantly influenced by sex or diet differences for gene and isoform abundance (FDR< 0.1; LOD ≥10.5). **(B)** The Venn diagrams show overlaps between the number of sex- and diet-influenced unique mRNAs corresponding to gene- and isoform QTL. **(C-D)** Distal QTL mapping profiles of sex- **(C)** and diet-influenced **(D)** gene- and isoform-level QTL and pathway enrichment (LOD ≥ 10.5; FDR < 0.1). The density plots show the number of distal QTL occurring within a 4Mbp window for distal gene- and isoform-level QTL (middle) and hotspot enrichments (top and bottom). **(E-F).** Isoform-level sex- and diet-specific distal QTL. Density plots show the number of QTL per 4 Mb window for unique mRNAs observed in either sex (**E**) or diet (**F**). **(G-H)** The QTL effect heatmaps highlight the cholesterol mRNAs that co-map in a sex- and diet-specific manner on chromosomes 7 and 9, respectively. **(I)** The Manhattan plots show local gene-level QTL (FDR < 0.1; LOD ≥ 10.5) associated with sex- and diet-specific mRNAs (occurring in one sex/diet group but not the other). The highlighted genes have LOD scores > 40 (sex) and 15 (diet). **(J)** The heatmap shows HuGE Scores of sex- and diet-specific genes associated with metabolic phenotypes in humans (genes highlighted on the plots depict those with the highest LOD scores for each sex and diet group detected for each trait).

**S7A Fig. (A)** Detection of sample mix-ups by QTL concordance. For each of the four molecular phenotypes, we selected the strongest local QTL. At each QTL peak marker, we predicted every mouse’s phenotype from its genotype as the product of the eight founder-allele dosages and the estimated founder-allele effects. Observed phenotypes were rank-normal transformed and adjusted for the covariates used in mapping (generation/litter, sex, diet), matching the QTL model. For every pair of mice (*i*, *j*), we then computed the Pearson correlation between the observed profile of mouse (*i*) and the genotype-predicted profile of mouse (*j*) across the selected QTL, yielding an N×N concordance matrix per molecular layer. Correctly labeled samples’ observed profiles are expected to correlate most strongly with the prediction from their own genotype; this is captured along the diagonal line. **(B)** PCA plots show batch effects on data for each molecular layer.

### Supplementary Tables

**S1 Table**. Gene-expression QTL (eQTL) (FDR < 0.1).

**S2 Table**. Isoform-abundance QTL (isoQTL) (FDR < 0.1).

**S3 Table**. Isoform-ratio QTL (irQTL) (FDR < 0.1).

**S4 Table**. Splice-junction QTL (sQTL) (FDR < 0.1).

**S5 Table**. GO biological process enrichment of Concordant isoQTL isoforms (Enrichr GO_Biological_Process_2025; *Adj. P* < 0.1, >= 5 genes).

**S6 Table**. GO biological process enrichment of Discordant isoQTL isoforms (Enrichr GO_Biological_Process_2025; *Adj. P* < 0.1, >= 5 genes).

**S7 Table**. GO biological process enrichment of Inverted isoQTL isoforms (Enrichr GO_Biological_Process_2025; *Adj. P* < 0.1, >= 5 genes).

**S8 Table**. GO biological process enrichment of Discordant irQTL (isoform-ratio) isoforms (Enrichr GO_Biological_Process_2025; *Adj. P* < 0.1, >= 5 genes).

**S9 Table**. GO biological-process enrichment of Inverted irQTL (isoform-ratio) isoforms (Enrichr GO_Biological_Process_2025; *Adj. P* < 0.1, >= 5 genes).

**S10 Table**. Gene-level enrichment of co-mapping distal QTL hotspots: GO biological-process terms per locus (*Adj. P* < 0.1, >= 5 genes); hotspot position is the mid peak- enrichment -+4-Mb bin.

**S11 Table**. Isoform-level enrichment of co-mapping distal QTL hotspots: GO biological-process terms per locus (*Adj. P* < 0.1, >= 5 genes); hotspot position is the mid peak- enrichment -+4-Mb bin.

**S12 Table**. Enrichment of isoform-unique distal QTL hotspots: GO biological-process terms per locus (*Adj. P* < 0.1, >= 5 genes); hotspot position is the mid peak-enrichment - +4-Mb bin.

**S13 Table**. GO biological-process enrichment of isoform-ratio (irQTL) distal-QTL hotspots: GO biological-process terms per locus (*Adj. P* < 0.1, >= 5 genes); hotspot position is the mid peak-enrichment -+4-Mb bin.

**S14 Table**. GO biological-process enrichment of splice-junction (sQTL) distal-QTL hotspots: GO biological-process terms per locus (*Adj. P* < 0.1, >= 5 genes); hotspot position is the mid peak-enrichment -+4-Mb bin.

**S15 Table**. Sex-dependent gene-expression QTL (QTLxSex eQTL) (FDR < 0.1).

**S16 Table**. Sex-dependent isoform-abundance QTL (QTLxSex isoQTL) (FDR < 0.1).

**S17 Table**. Diet-dependent gene-expression QTL (QTLxDiet eQTL) (FDR < 0.1).

**S18 Table**. Diet-dependent isoform-abundance QTL (QTLxDiet isoQTL) (FDR < 0.1).

**S19 Table**. GO biological process enrichment of sex-dependent gene-level distal QTL hotspots. Enriched terms per locus (*Adj. P* < 0.1, >= 5 genes); hotspot position is the mid peak-enrichment -+4-Mb bin.

**S20 Table**. GO biological-process enrichment of sex-dependent isoform-level distal- QTL hotspots. Enriched terms per locus (*Adj. P* < 0.1, >= 5 genes); hotspot position is the mid peak-enrichment -+4-Mb bin.

**S21 Table**. GO biological-process enrichment of diet-dependent gene-level distal-QTL hotspots. Enriched terms per locus (*Adj. P* < 0.1, >= 5 genes); hotspot position is the mid peak-enrichment -+4-Mb bin.

**S22 Table**. GO biological-process enrichment of diet-dependent isoform-level distal- QTL hotspots. Enriched terms per locus (*Adj. P* < 0.1, >= 5 genes); hotspot position is the mid peak-enrichment -+4-Mb bin.

**S23 Table**. Female-mouse gene-expression QTL (FDR < 0.1).

**S24 Table**. Male-mouse gene-expression QTL (FDR < 0.1).

**S25 Table**. HC-diet gene-expression QTL (FDR < 0.1).

**S26 Table**. HF-diet gene-expression QTL (FDR < 0.1).

**S27 Table**. Female-mouse isoform-abundance QTL (FDR < 0.1).

**S28 Table**. Male-mouse isoform-abundance QTL (FDR < 0.1).

**S29 Table**. HC-diet isoform-abundance QTL (FDR < 0.1).

**S30 Table**. HF-diet isoform-abundance QTL (FDR < 0.1).

## Notes

### Competing Interest Statement

The authors have declared no competing interest.

### Summary of Updates

This version details major revisions to the manuscript body, methodology, and results.

