## Supplementary figures and images for "Distinct genetic architectures of mRNA expression and splicing QTL in the Diversity Outbred mouse population"

### S1 Fig

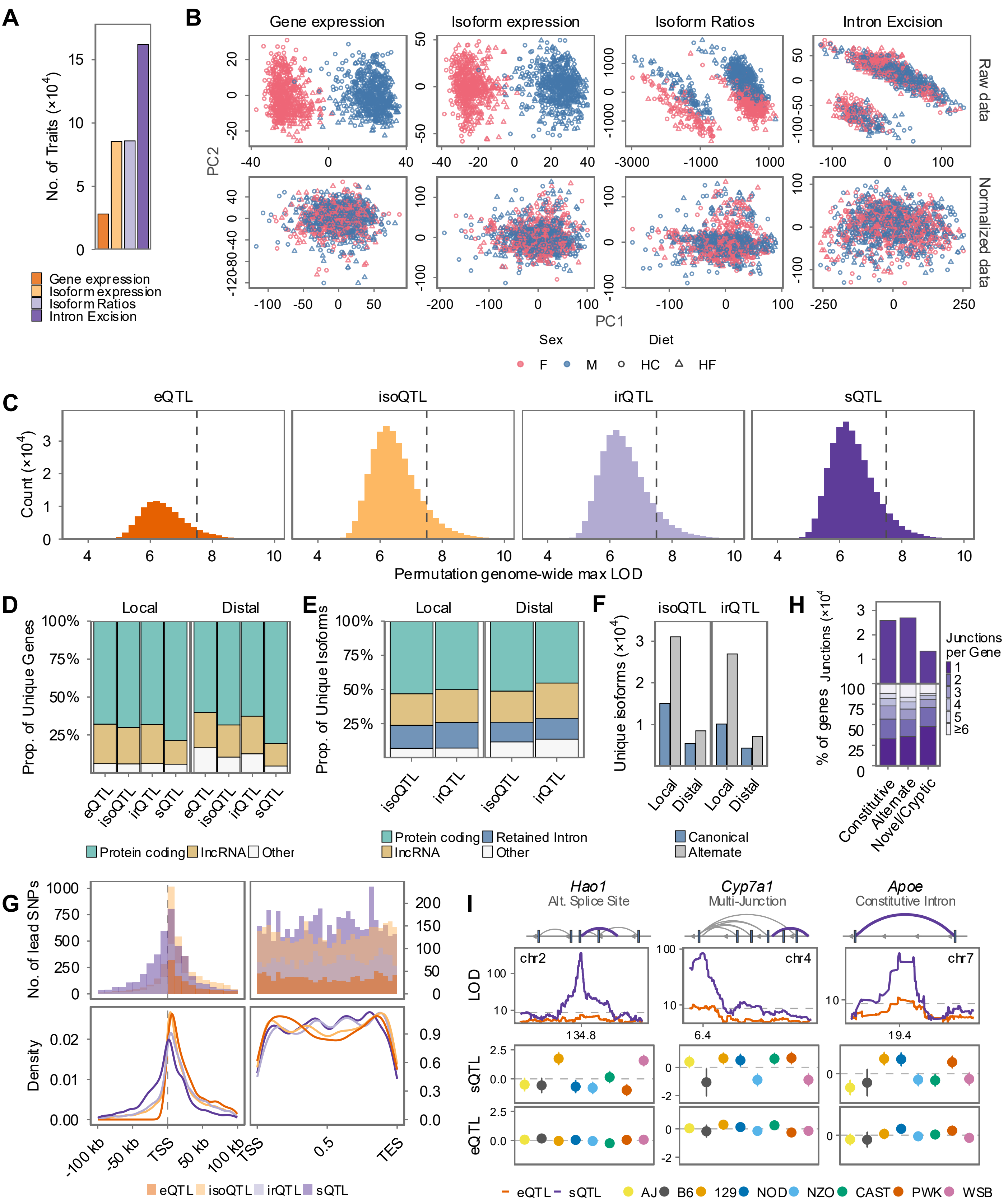

### S2 Fig

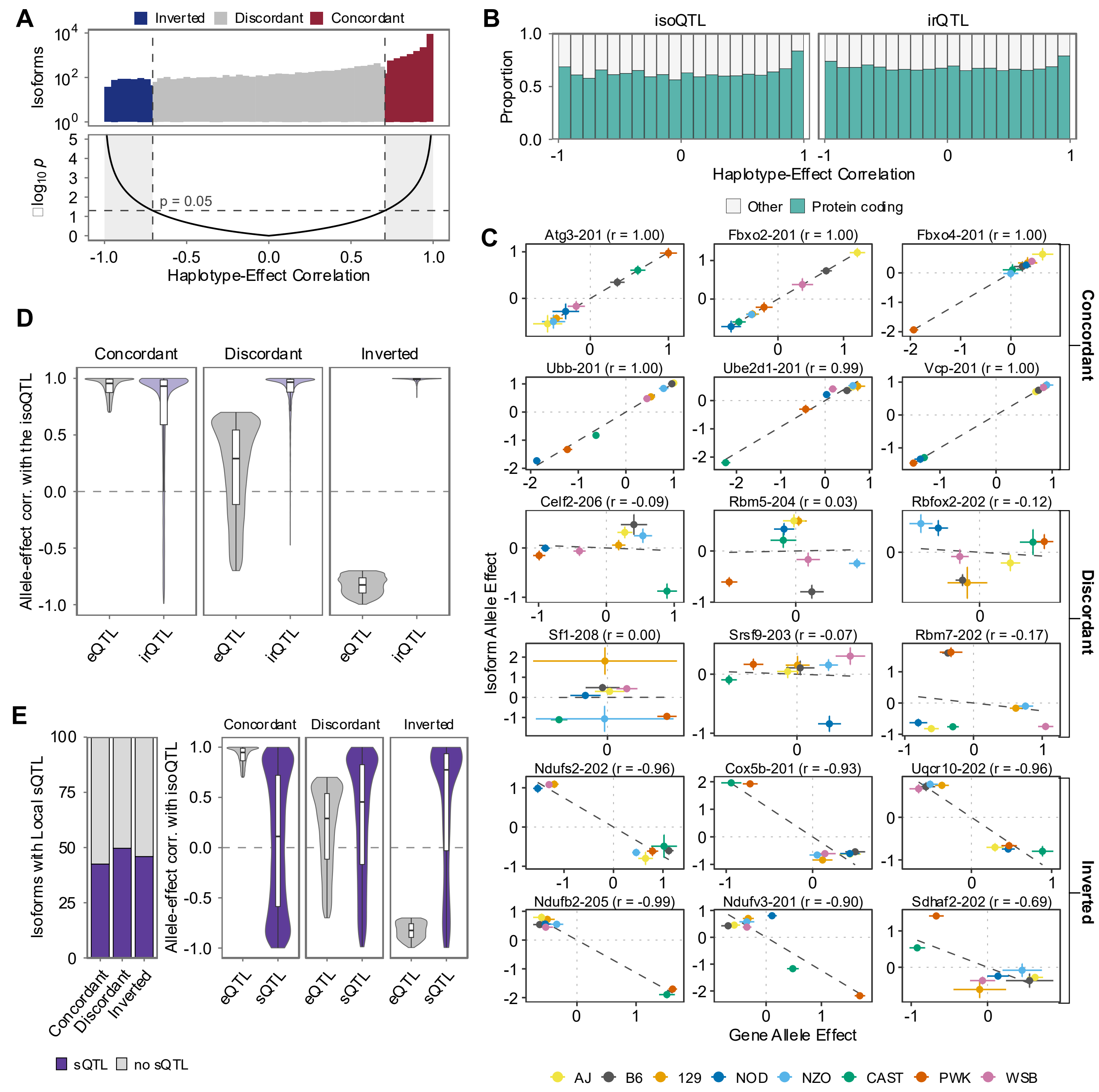

### S3 Fig

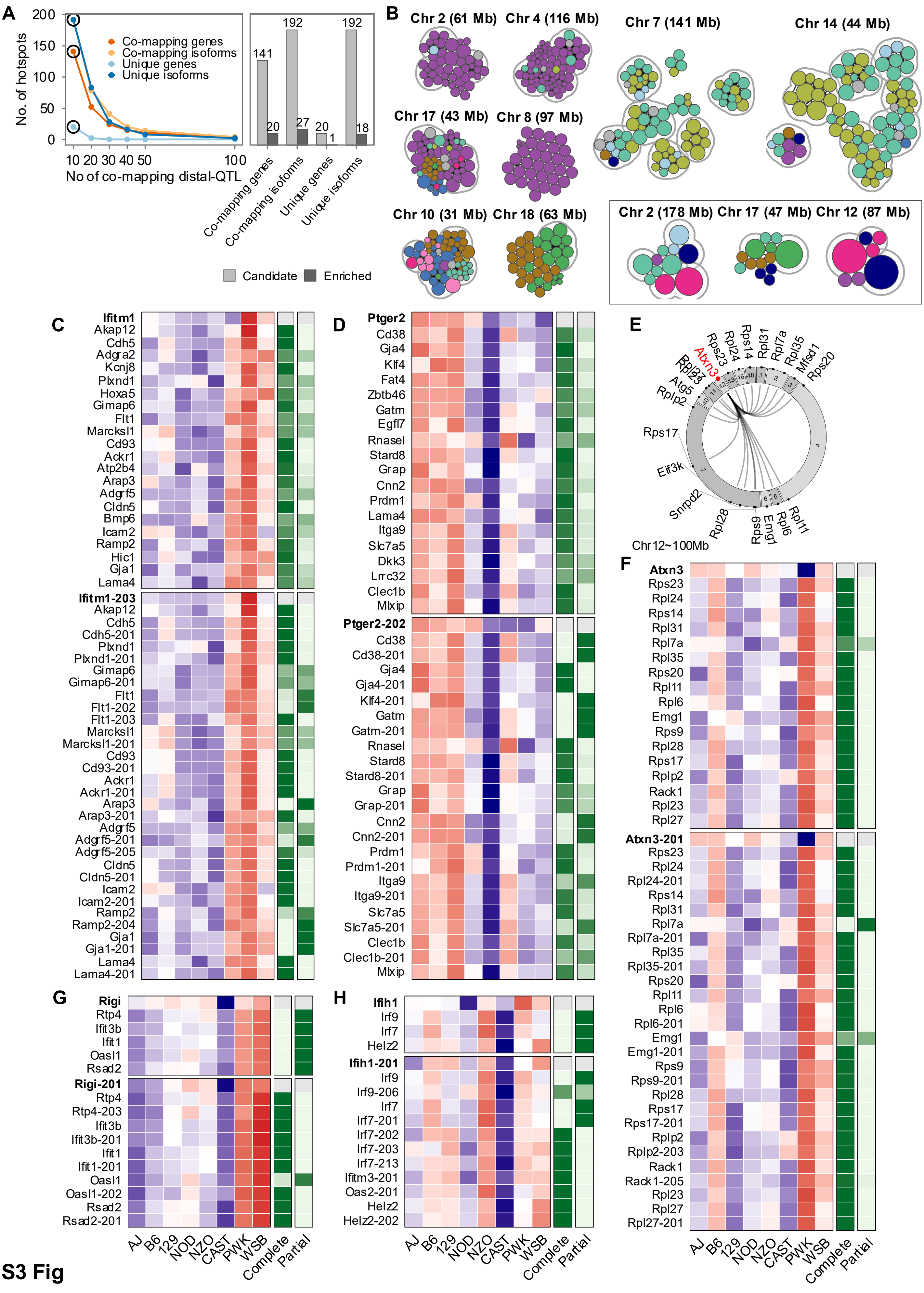

### S4 Fig

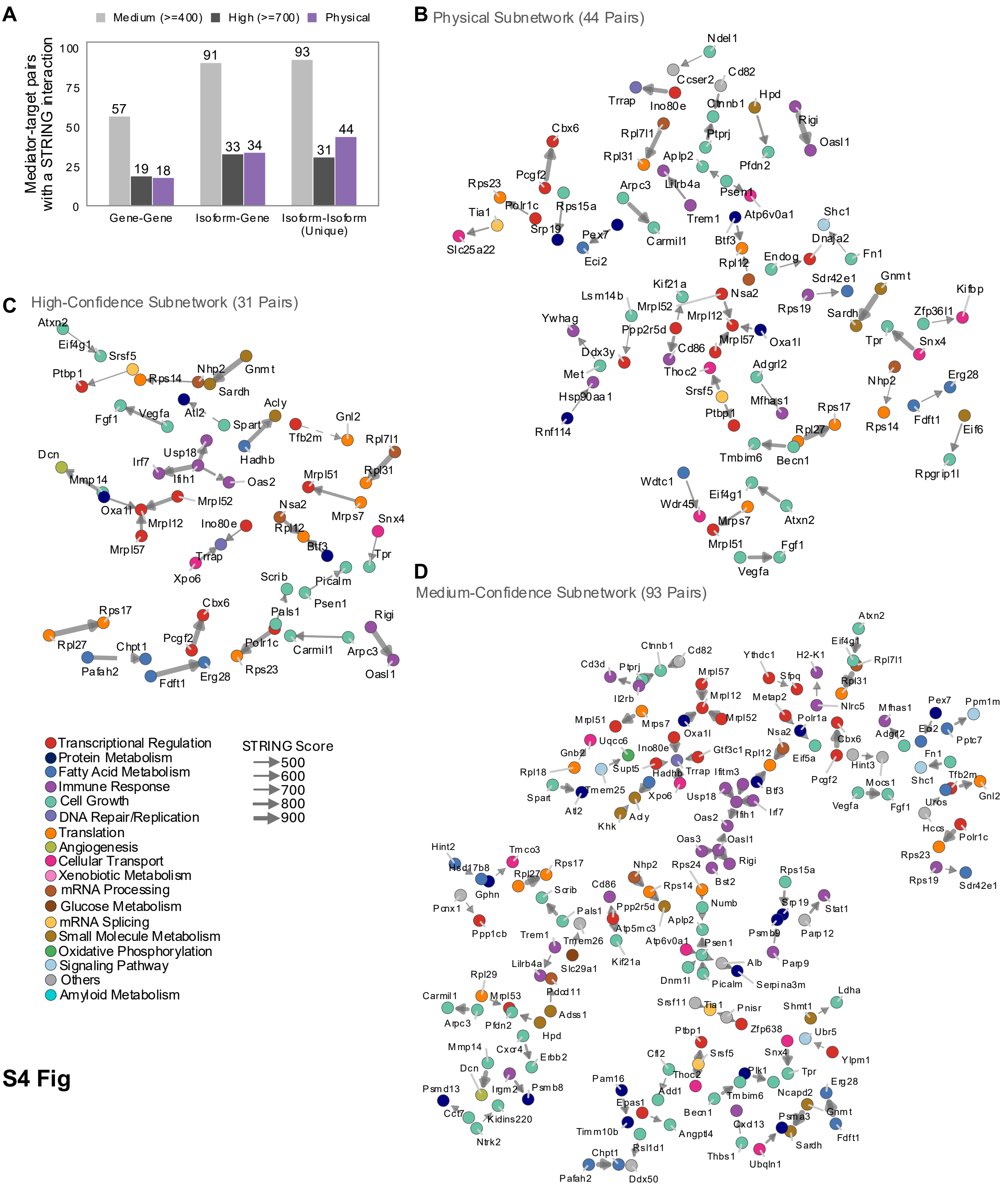

### S4 Fig

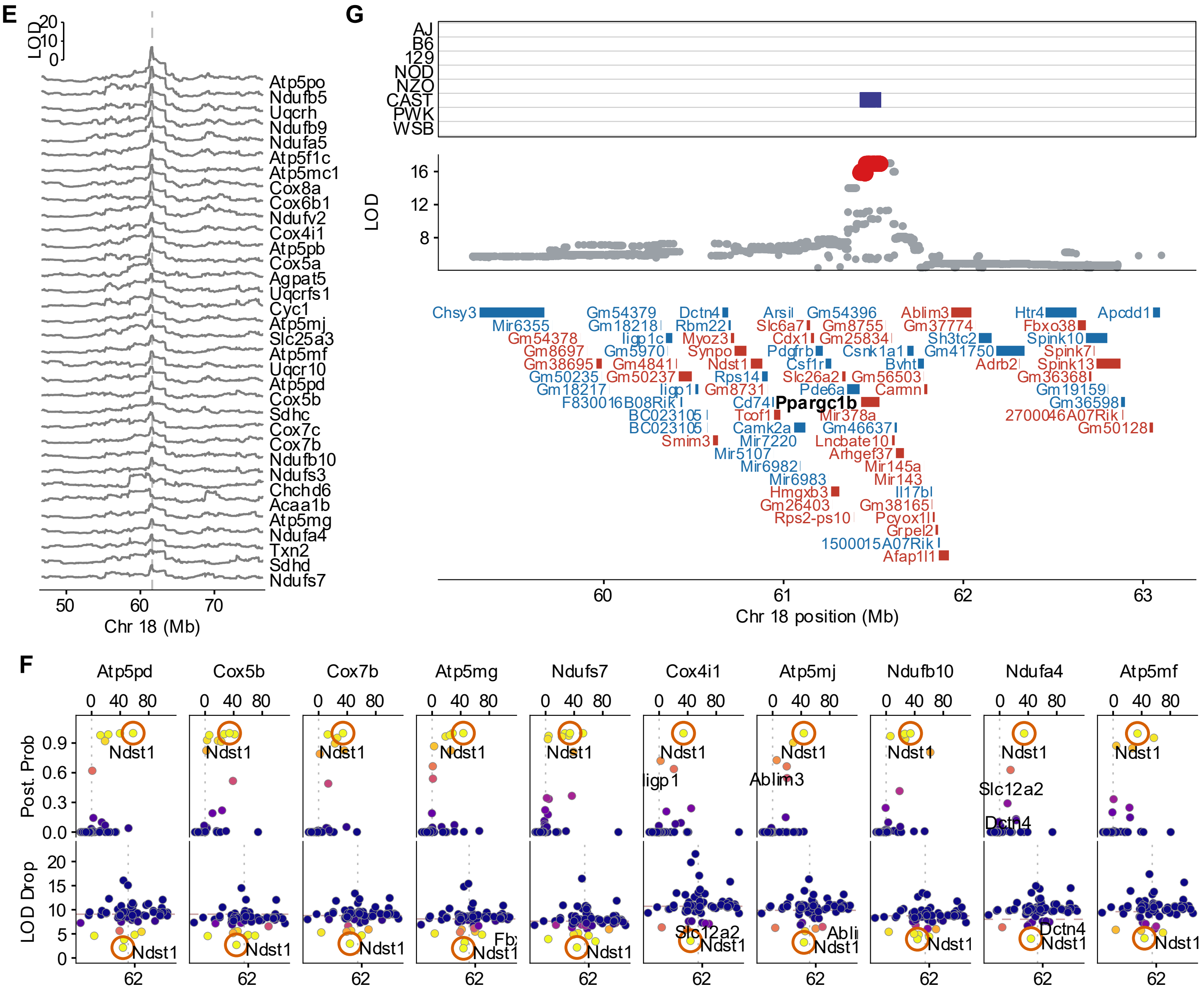

### S4 Fig

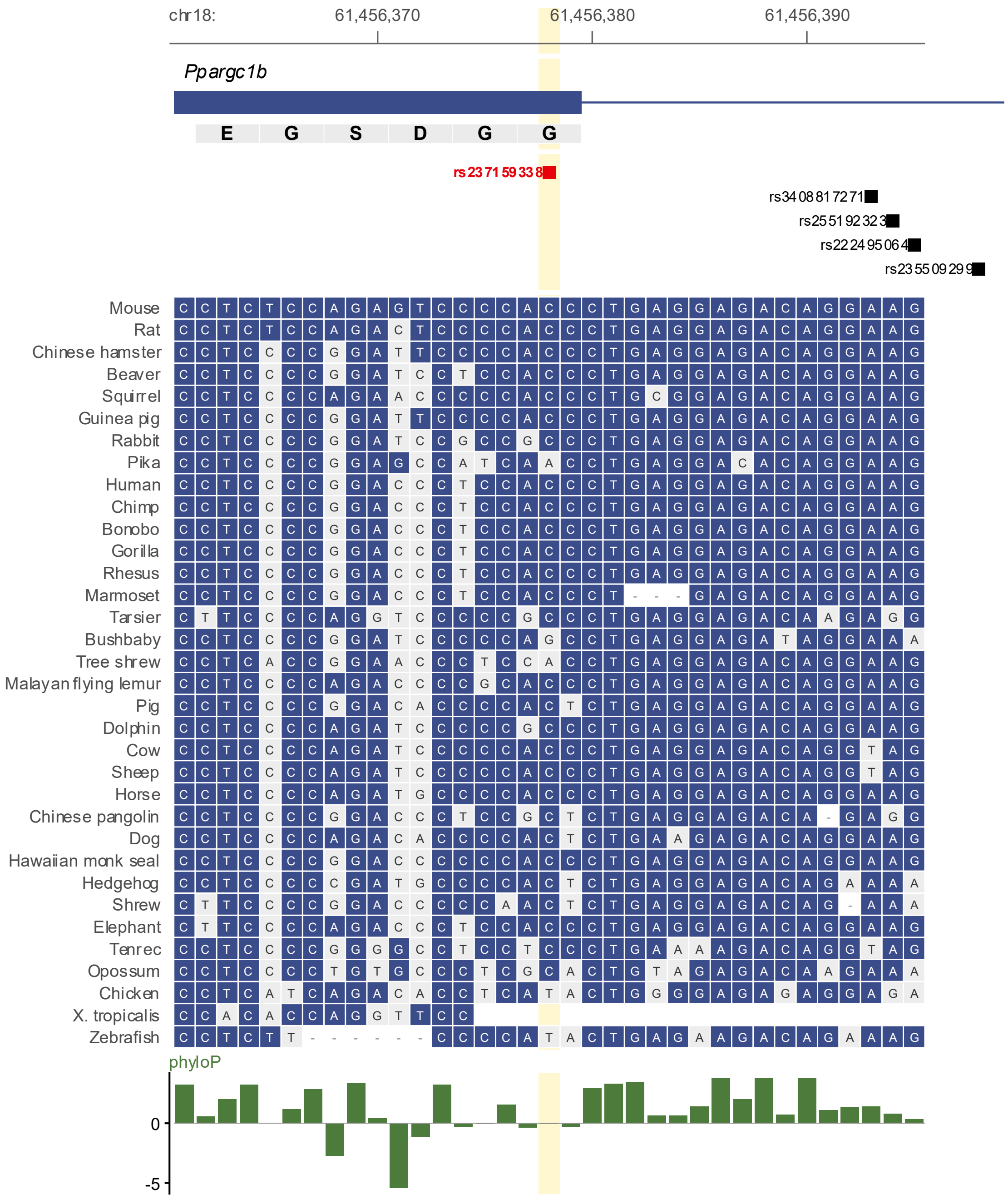

### S4 Fig

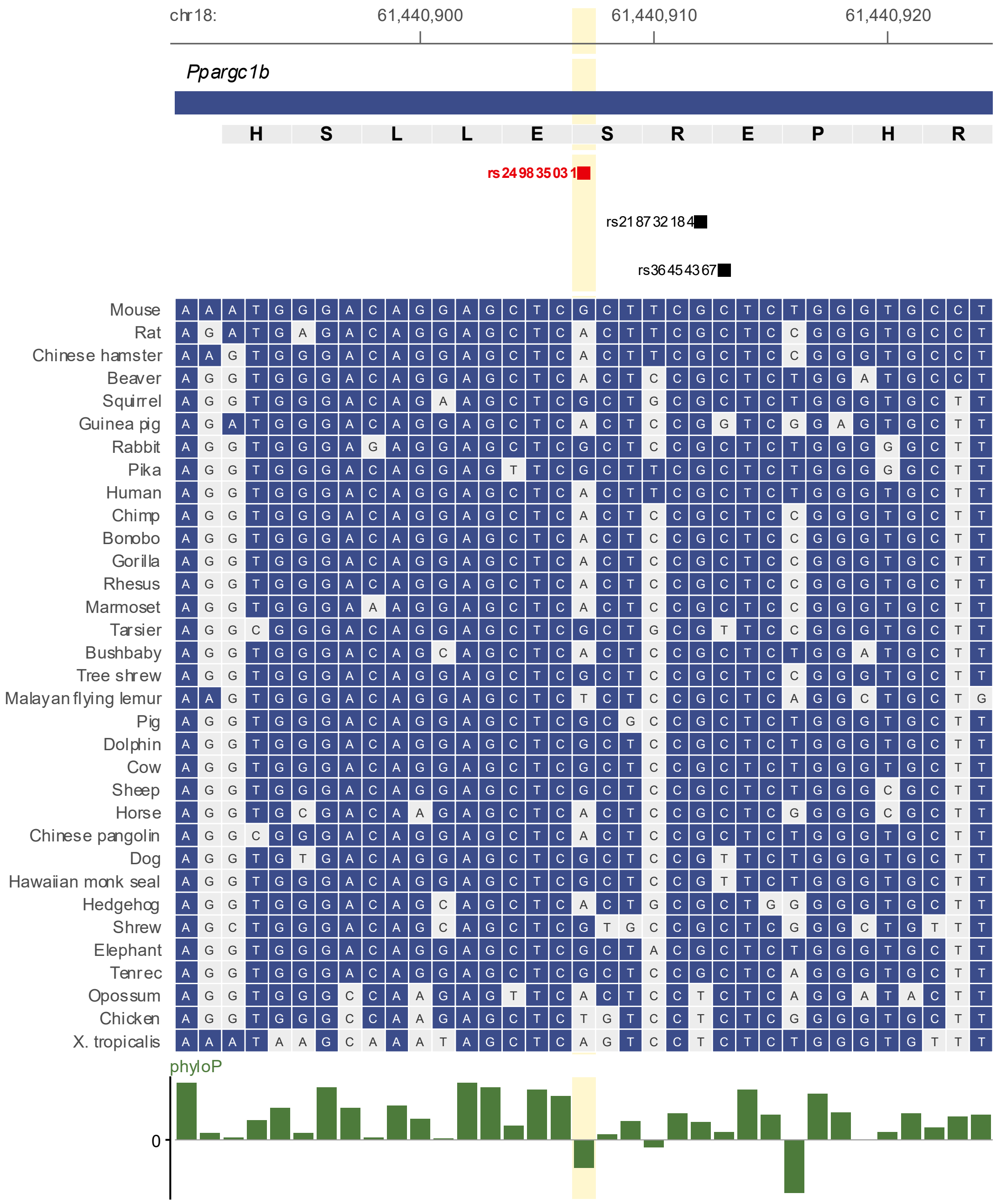

### S4 Fig

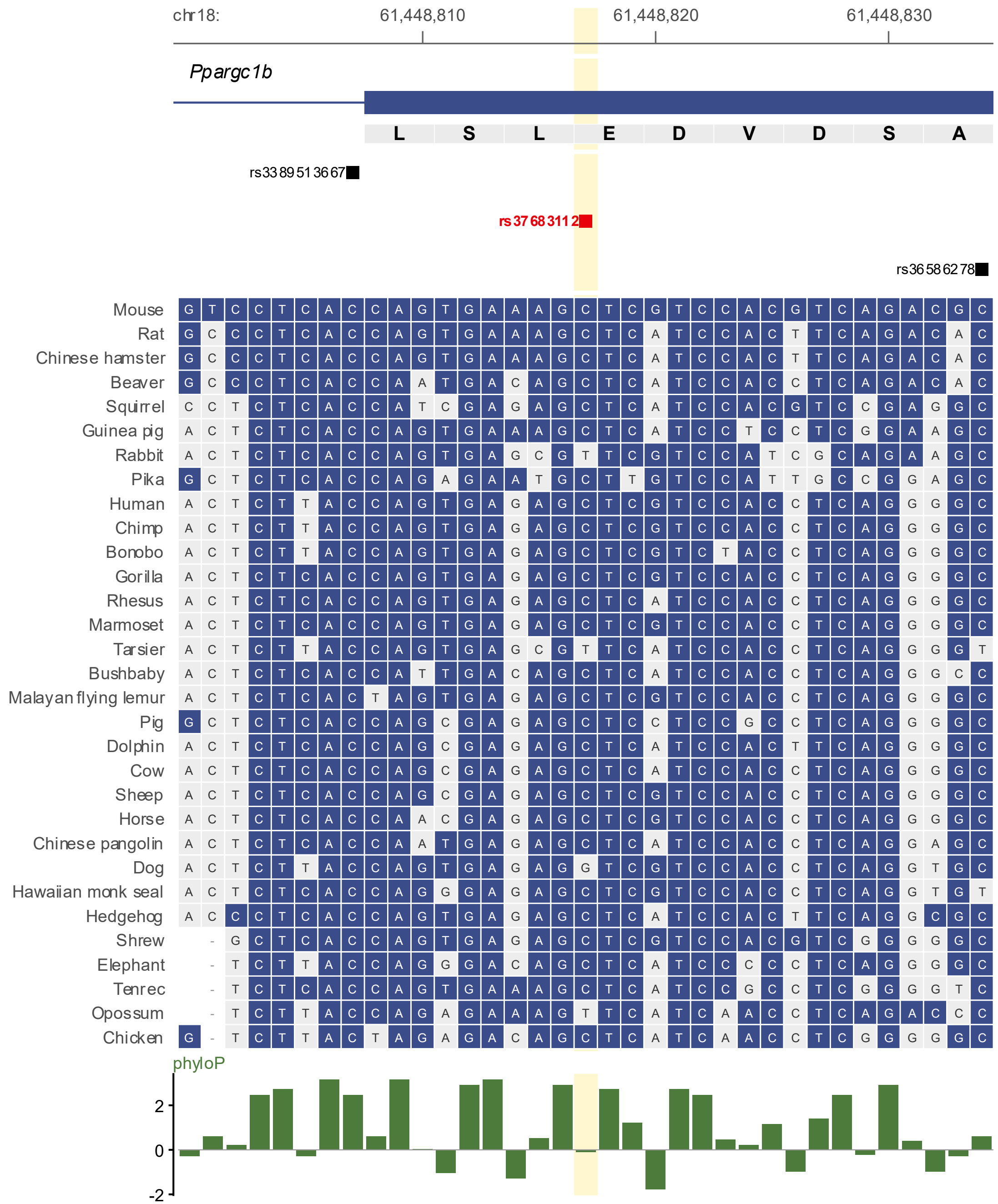

### S5 Fig

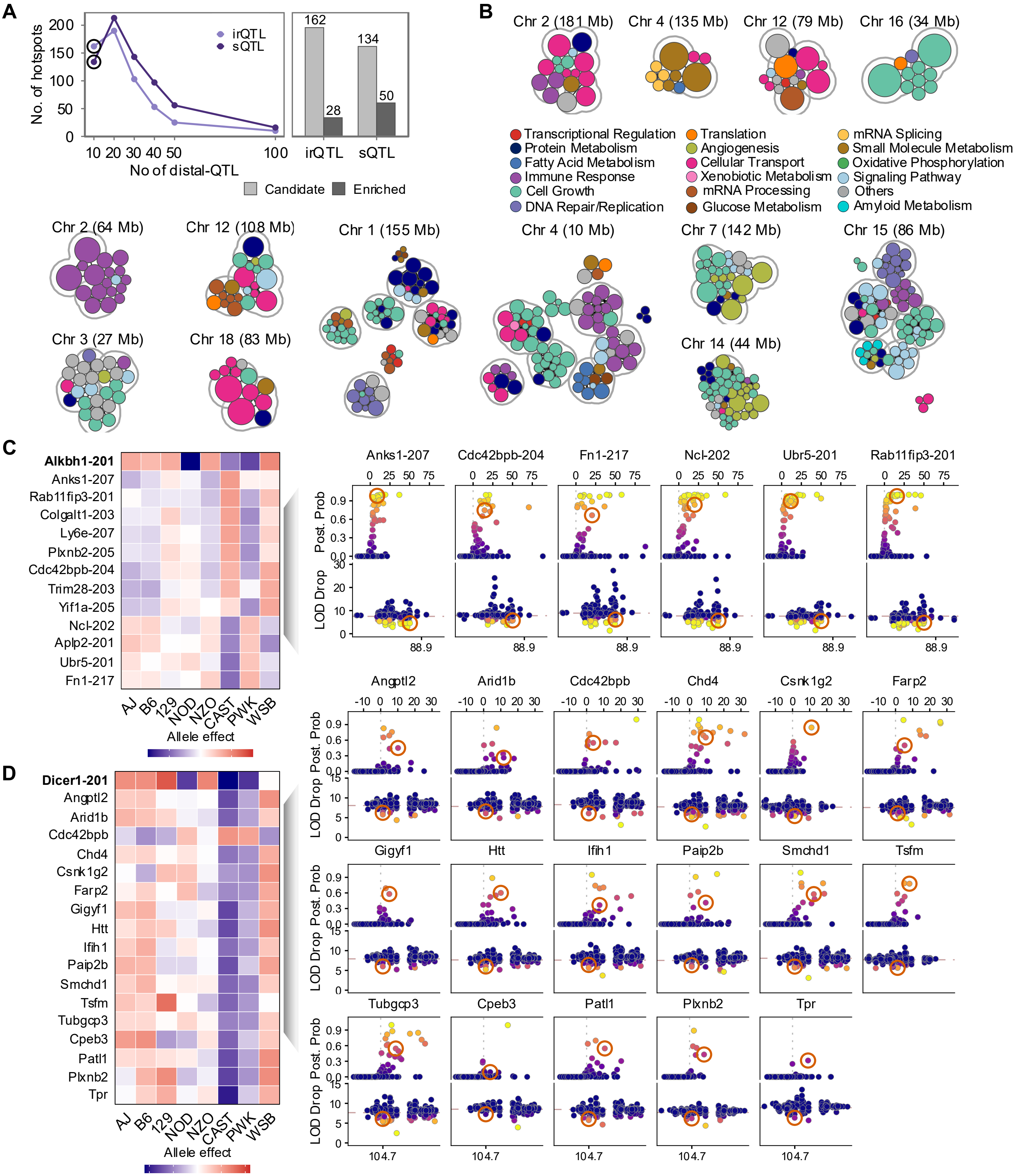

### S6 Fig

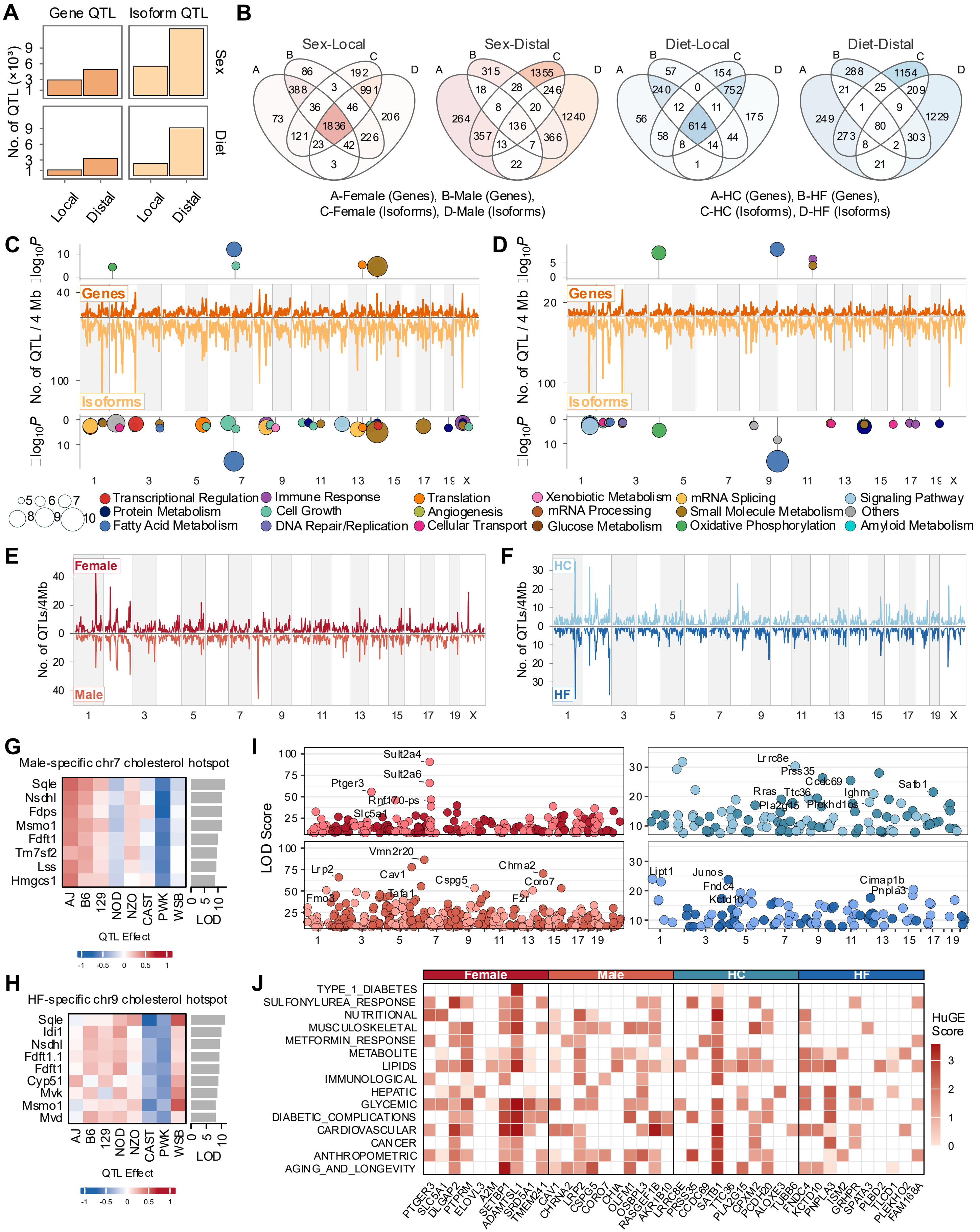

### S7 Fig

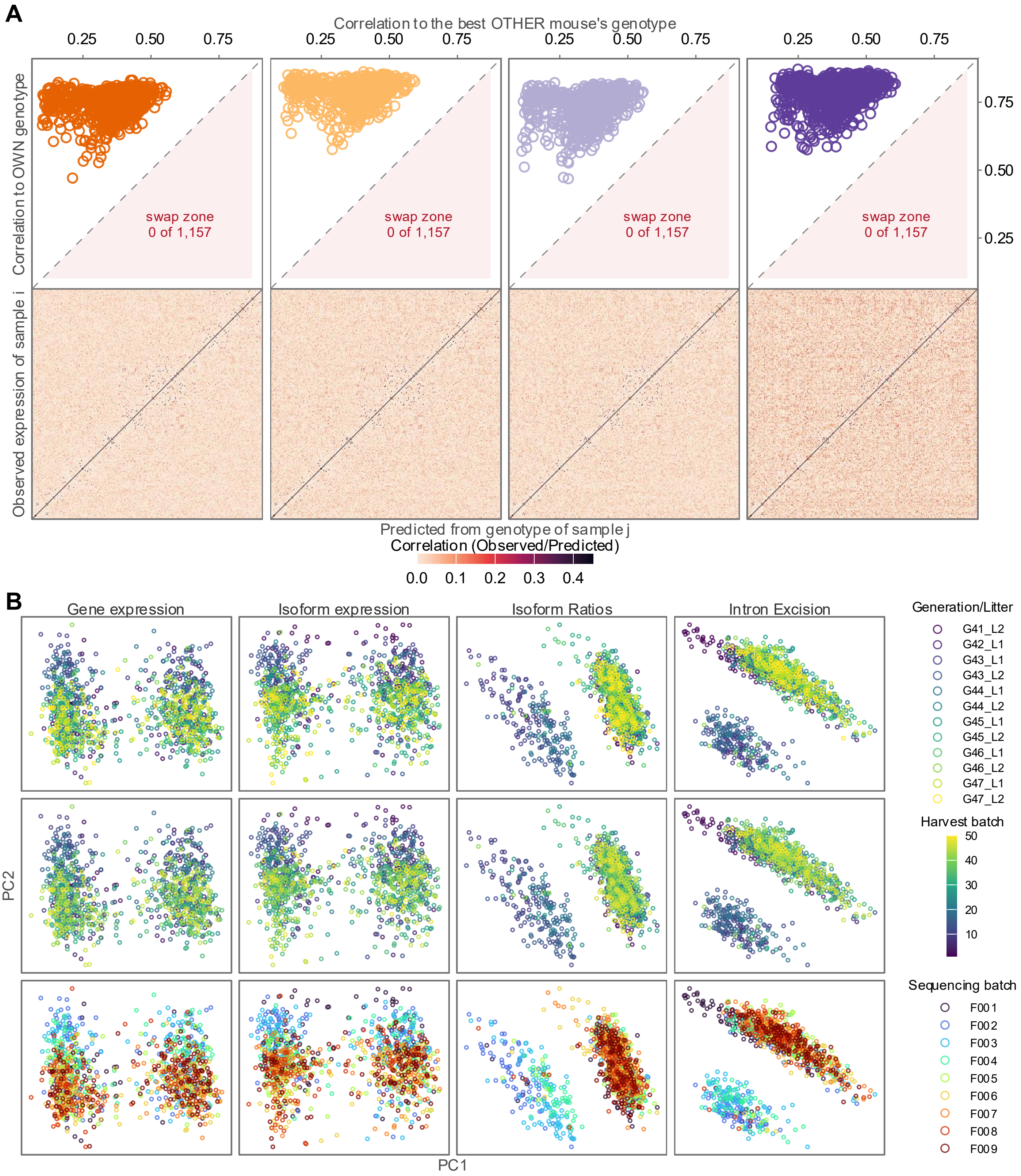
